# Dysregulated coordination of *MAPT* exon 2 and exon 10 splicing underlies different tau pathologies in PSP and AD

**DOI:** 10.1101/2021.09.23.461598

**Authors:** Kathryn R. Bowles, Derian A. Pugh, Laura-Maria Oja, Benjamin M. Jadow, Kurt Farrell, Kristen Whitney, Abhijeet Sharma, Jonathan D. Cherry, Towfique Raj, Ana C. Pereira, John F. Crary, Alison M. Goate

**Author notes:** Neuropathology Brain Bank & Research Core, Icahn School of Medicine at Mount Sinai, New York, NY, United States of America.

## Abstract

Understanding regulation of *MAPT* splicing is important to the etiology of many nerurodegenerative diseases, including Alzheimer disease (AD) and progressive supranuclear palsy (PSP), in which different tau isoforms accumulate in pathologic inclusions. *MAPT*, the gene encoding the tau protein, undergoes complex alternative pre-mRNA splicing to generate six isoforms. Tauopathies can be categorized by the presence of tau aggregates containing either 3 (3R) or 4 (4R) microtubule binding domain repeats (determined by inclusion/exclusion of exon 10), but the role of the N terminal domain of the protein, determined by inclusion/exclusion of exons 2 and 3 has been less well studied. Using an unbiased correlational screen in human brain tissue, we observed coordination of *MAPT* exons 2 and 10 splicing. Expression of exon 2 splicing regulators and subsequently exon 2 inclusion are differentially disrupted in PSP and AD brain, resulting in the accumulation of 1N4R isoforms in PSP and 0N isoforms in AD temporal cortex. Furthermore, we identified different N-terminal isoforms of tau present in neurofibrillary tangles, dystrophic neurites and tufted astrocytes, indicating a role for differential N-terminal splicing in the development of disparate tau neuropathologies. We conclude that N-terminal splicing and combinatorial regulation with exon 10 inclusion/exclusion is likely to be important to our understanding of tauopathies.

## INTRODUCTION

Ninety-five percent of all human multi-exonic genes are subject to alternative pre-mRNA splicing^1, 2^. Correct regulation of this mechanism is essential for proteomic diversity by the production of multiple distinct isoforms from a single gene^3^. The microtubule-associated protein tau (*MAPT*) is a neuronally expressed gene consisting of 16 exons, many of which are differentially spliced within the central nervous system and peripheral tissues. Tau proteins are involved in axonal transport, synaptic plasticity, and stabilization of the microtubule network^4–6^. In the human brain, the splicing of *MAPT* exons 2, 3 and 10 results in the expression of six different isoforms, which can be parsed into two groups depending on their inclusion or exclusion of exon 10 (Figure 1A).

**Figure 1.**
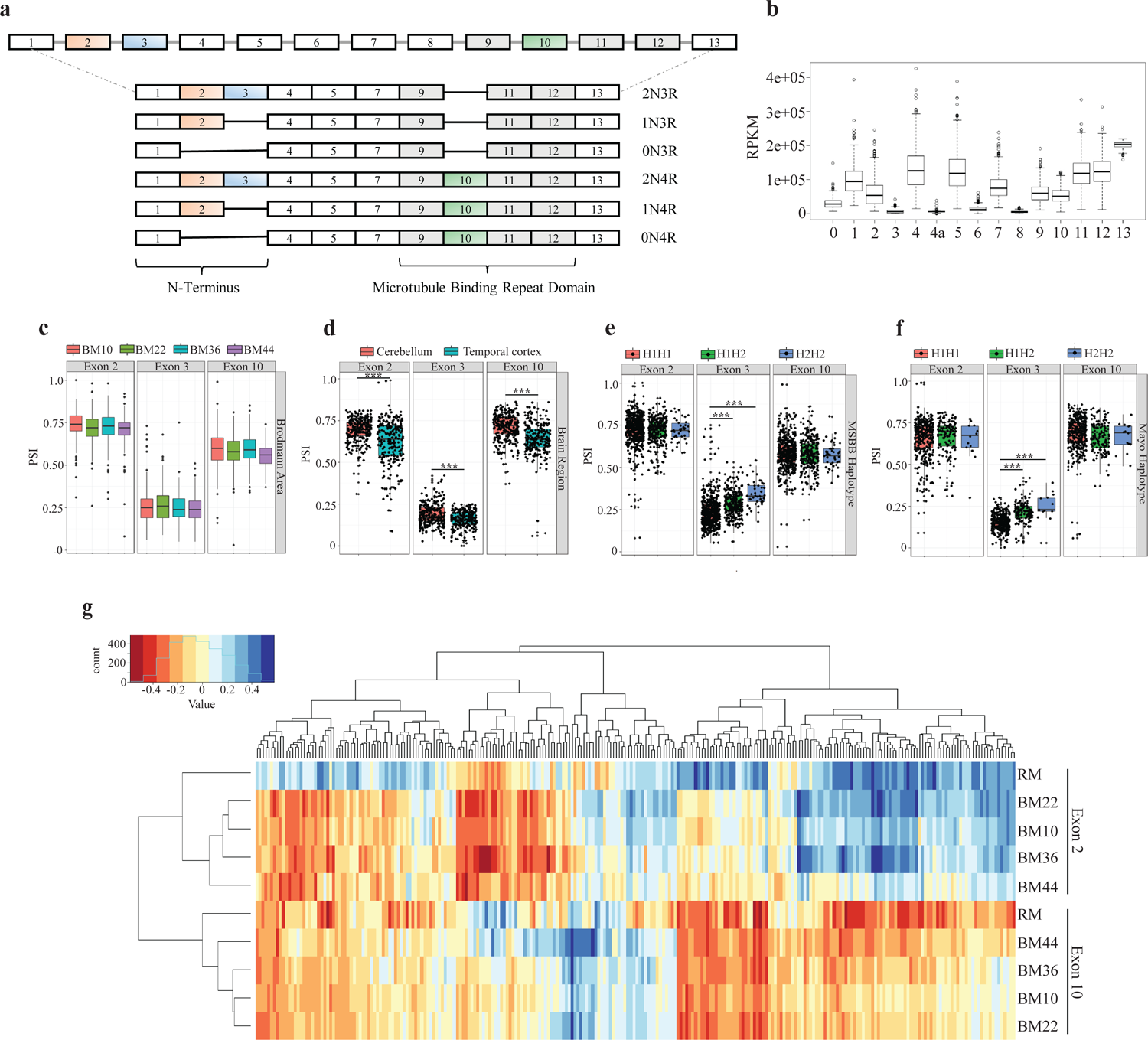
Splicing factor and RNA binding protein expression is differentially correlated with the inclusion of *MAPT* exons 2 and 10. **A)** *MAPT* exons 2, 3 and 10 are alternatively spliced, resulting in the expression of 6 different isoforms. At the N-terminus, exons 2 and 3 may be included or excluded, although exon 3 inclusion requires the inclusion of exon 2. The absence of either exon results in 0N isoforms, exon 2 alone results in 1N isoforms, and exon 3 inclusion defines 2N isoforms. At the C-terminus, the inclusion of exon 10 (encoding the second microtubule binding repeat domain) defines 4R isoforms, whereas its exclusion results in 3R isoforms. **B)** *MAPT* exon expression (RPKM) in the AMP-AD ROSMAP cohort. Error bars ± SEM. **C)** *MAPT* exons 2, 3 and 10 PSI across four Brodmann regions examined in the AMP-AD MSBB cohort. Error bars ± SEM. **D)** PSI values for *MAPT* exons 2, 3 and 10 in cerebellum and temporal cortex in the AMP-AD MAYO cohort. Error bars ± SEM. **E-F)** *MAPT* PSI values for exons 2, 3 and 10 between 17q21.31 H1 and H2 haplotypes in ***E.*** MSBB and ***F.*** MAYO AMP-AD datasets. Error bars ± SEM. **G)** Pearson’s correlation coefficients with unsupervised hierarchical clustering between SF/RBP expression (x-axis) and *MAPT* exon 2/exon 10 PSI values in ROSMAP and MSBB data. Blue indicates a positive correlation (“includers”) and red indicates a negative correlation (“excluders”), while yellow denotes no association. All comparisons carried out using linear regression model. \*\*\**p* < 0.001

The ratio of tau isoform expression changes during human brain development^7, 8^, with only the shortest 0N3R isoform of tau being expressed in fetal brain^7–10^. Following birth, there is a sudden shift in the expression of both exons 2 and 10; exon 10 inclusion increases dramatically during the perinatal period^8^ where it reaches a stable 3R:4R ratio of roughly 1:1^11–13^, whereas exon 2 expression increases gradually throughout the first decade of life^8^. The reason for this shift and the function of different *MAPT* isoforms is not fully understood. However, 4R tau has an increased affinity for binding microtubules that results in their increased stabilization^14^, therefore shorter 3R isoforms may allow for greater neuroplasticity during brain development. It has been proposed that different N-terminal isoforms may also contribute to microtubule stabilization^15^, and that the inclusion of exons 2 and 3 results in the extension of the acidic region of tau, lengthening its projection domain^16^, which in turn may increase the distance between microtubules and increase bundling^17^.

Understanding *MAPT* splicing is of critical importance to the etiology of tauopathies, which are characterized by the presence of neuronal and/or astroglial tau aggregates. Primary tauopathies include frontotemporal dementia (FTD), progressive supranuclear palsy (PSP), corticobasal degeneration (CBD), primary age-related tauopathy (PART) and Pick’s disease (PiD), while other tauopathies, such as AD, are secondary to amyloid-beta (Aß) deposition. In several primary tauopathies the regulation of *MAPT* splicing is altered and mis-spliced isoforms are differentially incorporated into neurofibrillary tangles (NFTs) and pathogenic inclusions^18, 19^. The most striking evidence in support of the importance of regulated splicing is the numerous synonymous and intronic *MAPT* mutations, such as S305S^20, 21^, IVS10+16^22^ and N296N^23^, which result in increased exon 10 inclusion, and the subsequent development of autosomal dominant FTD. The ability for these mutations to induce tau pathology in the absence of an altered amino acid sequence is indicative of the relevance of *MAPT* splicing to disease pathogenesis, and the importance of maintaining the correct tau isoform ratio.

Alternative splicing of *MAPT* exon 10 in both healthy and diseased brain has been well characterized, although studies examining exon 10 expression in AD have yielded inconsistent results^11, 13, 24–27^. In contrast, less is known about the regulation of exons 2 and 3, and there have been no studies directly assessing the contribution of N-terminal tau isoforms to primary tauopathies. To date, splicing of exon 10 has been associated with the function of several candidate splicing factors (SFs) and RNA binding proteins (RBPs), the most comprehensively investigated of which are the serine and arginine-rich family of splicing factors (SRSFs). Multiple SRSFs have been associated with both exon 10 inclusion and exclusion^28–31^, as have Tra2β^19, 29, 32, 33^, FUS^33^, RBM4^34^, NOVA1^35^ and hnRNPs E2 and E3^35, 36^. However, there is limited evidence for many of these associations, and their impact on *MAPT* splicing lacks robust replication.

Here, we report a correlational screen for SFs/RBPs in human brain tissues that revealed novel genes associated with *MAPT* splicing, and uncovered coordinated regulation of *MAPT* exons 2 and 10. We found that the splicing factor *RSRC1* bound directly to *MAPT* pre-mRNA and was associated with both exon 10 exclusion and exon 2 inclusion. Furthermore, *RSRC1* was also differentially expressed in PSP and AD brain, suggesting a regulatory role for this SF in disease pathogenesis. Consistent with discordant *RSRC1* expression, we observed increased expression of exon 2 and exon 10-containing transcripts in PSP brain, and increased expression of 0N transcripts in AD brain, which correlate with the accumulation of different N-terminal tau isoforms in different neuropathological features of AD and PSP. We therefore conclude that differential expression of *MAPT* splicing regulators in AD and PSP brain results in the loss of coordination between *MAPT* exon 10 and exon 2 splicing. In turn, this results in the expression and accumulation of different N-terminal isoforms in each disease, which may underlie the development of different neuropathological features characteristic of AD and PSP.

## RESULTS

### *MAPT* alternative splicing differs by brain region and 17q21.31 haplotype

*MAPT* is alternatively spliced at exons 2, 3 and 10 in the human brain (Figure 1A), and expression of these exons across brain regions has previously been measured by microarray analyses^12^. However, accurate measurement of gene or exon expression by microarray may be impacted by the specificity of probe design and a narrow dynamic range for signal detection. We therefore chose to characterize the relative expression of *MAPT* exons in multiple human postmortem RNA-seq datasets from the AMP-AD consortium (Religious Orders Study Memory and Aging Project [ROSMAP; Synapse syn3219045] N = 450, Mount Sinai Brain Bank [MSBB; Synapse syn3159438] N = 230, and the Mayo Clinic [MAYO; Synapse syn3157268, syn5550404] N = 276) (Figure 1B-F, Figure S1A-H). We observed a pattern of exon expression consistent with that previously described^12, 37^ and known isoforms expressed in brain (i.e., constitutive expression of exons 1, 4, 5, 7, 9, 11 and 12, little to no expression of exons 4a, 6 and 8, and variable levels of exon 2, 3 and 10), which was consistent across AMP-AD datasets and brain regions (Figure 1B, Figure S1A-C).

To assess proportional exon expression across different brain regions included in MSBB and MAYO data, we calculated percent spliced in (PSI) values for the alternatively spliced exons 2, 3 and 10 using MISO (Mixture of Isoforms) (Figure 1C-D). While there were no differences in PSI values across MSBB Brodmann regions BM10 (frontal pole), BM22 (superior temporal gyrus), BM36 (fusiform gyrus) and BM44 (inferior frontal gyrus; Figure 1C), there was significantly increased inclusion of all three exons in cerebellum compared to the temporal cortex in the MAYO cohort (Exon 2 *p* = 5.46×10^-16^; Exon 3 *p* = 2.74×10^-8^; Exon 10 *p* = 1.54×10^-9^; Figure 1D). This is consistent with previous reports that suggest *MAPT* splicing in the cerebellum differs from the forebrain^11, 12^.

We then compared PSI values between the major 17q21.31 *MAPT* H1 and H2 haplotypes, and observed increased exon 3 inclusion in H2 haplotype carriers compared to H1 across most AMP-AD datasets and brain regions (MSBB H1H2 *p*=1.14×10^-09^, MSBB H2H2 *p*=1.09×10^-13^; MAYO H1H2 *p*< 2×10^-16^, MAYO H2H2 *p*=1.9×10^-04^, Figure 1E-F, Figure S1D-E), which has been previously reported in other RNA-seq and microarray datasets^12, 38, 39^. In comparison, we did not observe any difference in exon 2 or 10 inclusion between haplotypes (Figure 1E-F, Figure S1D-E). In contrast to previous reports^40^, we did not find altered total *MAPT* expression between these haplotypes in any dataset or brain region (Figure S1F-H).

### *MAPT* exon 2 and 10 splicing is coordinated by SF/RBP expression

In order to identify novel *MAPT* splicing regulators in human prefrontal cortex, we carried out a correlational analysis between the expression (RPKM) of 294 known splicing factors (SFs) and RNA binding proteins (RBP) described in Gerstberger et al 2014^41^ (Table S1) with *MAPT* exon 2, 3 and 10 PSI values, using ROSMAP and MSBB datasets (Figure 1G, Figure S2A-G). We did not observe any significant correlations that withstood Bonferroni multiple test correction between *MAPT* exon 3 and any SF/RBP (Figure S2A), likely due to the very low expression of exon 3 in human brain. However, when we separated the data by H1/H2 haplotype, we observed stronger associations between exon 3 inclusion and SF/RBP expression in H2 homozygotes compared to H1 homozygotes in both datasets (Figure S2B-C), although these still did not pass multiple test correction due to low frequency of the H2 haplotype in these datasets. While we did not have the statistical power to pursue analysis of exon 3 splicing regulators, these data indicate there may be additional regulation of exon 3 in the context of H2, consistent with its increased expression on this background.

In contrast, there were no significant differences in SF/RBP and *MAPT* exon 2/10 PSI value associations between H1/H2 haplotypes, which were highly correlated across both haplotypes (ROSMAP exon 2 R^2^ = 0.81, *p* < 0.001, exon 10 R^2^ = 0.89, *p* < 0.001; MSBB exon 2 R^2^ = 0.67 *p* < 0.001, exon 10 R^2^ = 0.66 *p* < 0.001) (Figure S2D-E). Associations between PSI values and SF/RBP expression were also significantly correlated between AD cases and controls (ROSMAP exon 2 R^2^ = 0.97 *p* < 2.2×10^-16^, exon 10 R^2^ = 0.96 *p* < 2.2×10^-16^; MSBB exon 2 R^2^ = 0.89 *p* < 2.2×10^-16^, exon 10 R^2^ = 0.93 *p* < 2.2×10^-16^) (Figure S2F-G), indicating no differences in the regulatory effects of SF/RBP expression on *MAPT* splicing in AD. We therefore focused our analyses on exons 2 and 10 using pooled AD/control and H1/H2 data.

Unsupervised hierarchical clustering of the resulting Pearson’s correlation coefficients between exon PSI values and SF/RBP expression revealed that exons 2 and 10 clustered separately from each other and had distinct patterns of association with SF/RBP expression (Figure 1G). In order to identify robust *MAPT* splicing regulator candidates, we selected SF/RBPs with significant (Bonferroni-corrected *p*-value < 0.05) associations with *MAPT* exons 2 or 10 in the same direction across the three most anatomically similar datasets; ROSMAP (prefrontal cortex), MSBB BM10 (frontal pole) and MSBB BM44 (inferior frontal gyrus). Many more SFs/RBPs were associated with exon 10 exclusion (94 genes, 69.1% of all significant and replicated correlations, defined by a negative correlation; Table S2) compared with its inclusion (5 genes replicated in 2/3 datasets, [3.7%], defined by a positive correlation; Table S2), suggesting that more complex regulation may be required to promote removal of exon 10 from pre-mRNA transcripts in brain.

Fewer SFs/RBPs were associated with exon 2 splicing (7 [5.1%] excluders, 16 [11.8%] includers). However, a proportion of SFs/RBPs were significantly associated with both exon 2 and exon 10 PSI values in opposing directions (Table 1). Specifically, 14 SF/RBPs (10.3%) were significantly correlated with both exon 10 exclusion and exon 2 inclusion, suggesting a coordinated regulation of *MAPT* N- and C-terminal splicing that has not been previously characterized. We therefore chose to focus on this subset of genes (Table 1).

**Table 1.**
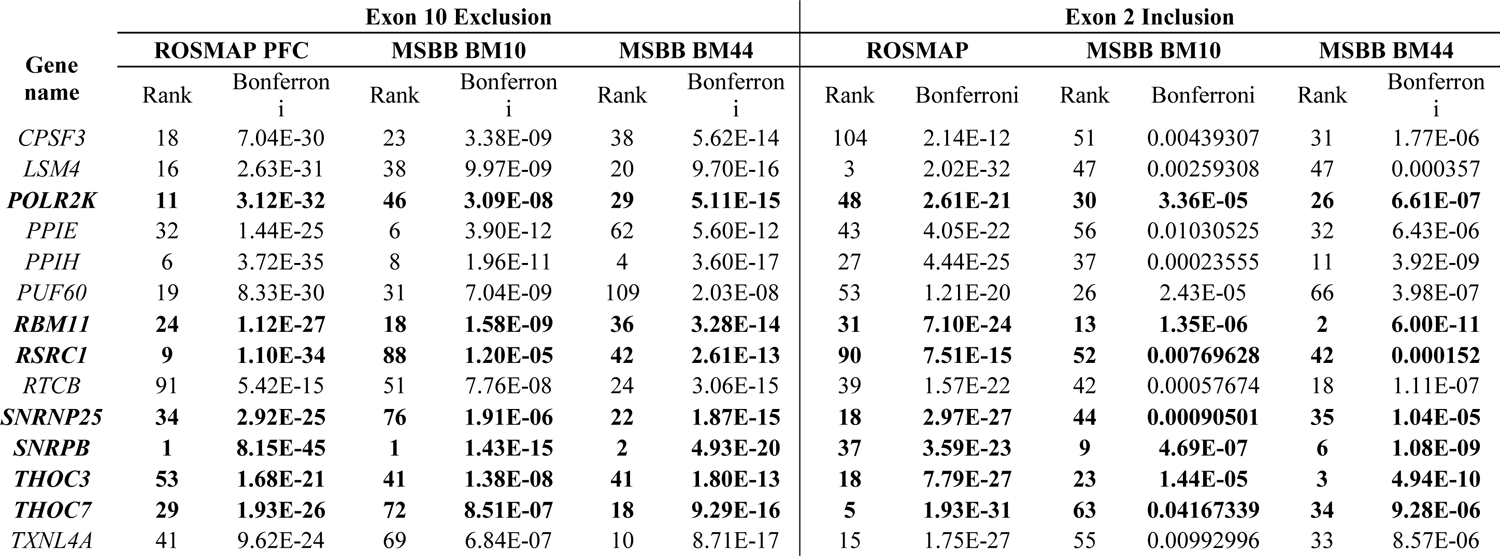
*MAPT* exon 10 excluders and exon 2 includers replicated across ROSMAP and MSBB datasets

| Gene name | Exon 10 Exclusion |  |  |  |  |  | Exon 2 Inclusion |  |  |  |  |  |
| --- | --- | --- | --- | --- | --- | --- | --- | --- | --- | --- | --- | --- |
|  | ROSMAP PFC |  | MSBB BM10 |  | MSBB BM44 |  | ROSMAP |  | MSBB BM10 |  | MSBB BM44 |  |
|  | Rank | Bonferroni | Rank | Bonferroni | Rank | Bonferroni | Rank | Bonferroni | Rank | Bonferroni | Rank | Bonferroni |
| *CPSF3* | 18 | 7.04E-30 | 23 | 3.38E-09 | 38 | 5.62E-14 | 104 | 2.14E-12 | 51 | 0.00439307 | 31 | 1.77E-06 |
| *LSM4* | 16 | 2.63E-31 | 38 | 9.97E-09 | 20 | 9.70E-16 | 3 | 2.02E-32 | 47 | 0.00259308 | 47 | 0.000357 |
| ***POLR2K*** | **11** | **3.12E-32** | **46** | **3.09E-08** | **29** | **5.11E-15** | **48** | **2.61E-21** | **30** | **3.36E-05** | **26** | **6.61E-07** |
| *PPIE* | 32 | 1.44E-25 | 6 | 3.90E-12 | 62 | 5.60E-12 | 43 | 4.05E-22 | 56 | 0.01030525 | 32 | 6.43E-06 |
| *PPIH* | 6 | 3.72E-35 | 8 | 1.96E-11 | 4 | 3.60E-17 | 27 | 4.44E-25 | 37 | 0.00023555 | 11 | 3.92E-09 |
| *PUF60* | 19 | 8.33E-30 | 31 | 7.04E-09 | 109 | 2.03E-08 | 53 | 1.21E-20 | 26 | 2.43E-05 | 66 | 3.98E-07 |
| ***RBM11*** | **24** | **1.12E-27** | **18** | **1.58E-09** | **36** | **3.28E-14** | **31** | **7.10E-24** | **13** | **1.35E-06** | **2** | **6.00E-11** |
| ***RSRC1*** | **9** | **1.10E-34** | **88** | **1.20E-05** | **42** | **2.61E-13** | **90** | **7.51E-15** | **52** | **0.00769628** | **42** | **0.000152** |
| *RTCB* | 91 | 5.42E-15 | 51 | 7.76E-08 | 24 | 3.06E-15 | 39 | 1.57E-22 | 42 | 0.00057674 | 18 | 1.11E-07 |
| ***SNRNP25*** | **34** | **2.92E-25** | **76** | **1.91E-06** | **22** | **1.87E-15** | **18** | **2.97E-27** | **44** | **0.00090501** | **35** | **1.04E-05** |
| ***SNRPB*** | **1** | **8.15E-45** | **1** | **1.43E-15** | **2** | **4.93E-20** | **37** | **3.59E-23** | **9** | **4.69E-07** | **6** | **1.08E-09** |
| ***THOC3*** | **53** | **1.68E-21** | **41** | **1.38E-08** | **41** | **1.80E-13** | **18** | **7.79E-27** | **23** | **1.44E-05** | **3** | **4.94E-10** |
| ***THOC7*** | **29** | **1.93E-26** | **72** | **8.51E-07** | **18** | **9.29E-16** | **5** | **1.93E-31** | **63** | **0.04167339** | **34** | **9.28E-06** |
| *TXNL4A* | 41 | 9.62E-24 | 69 | 6.84E-07 | 10 | 8.71E-17 | 15 | 1.75E-27 | 55 | 0.00992996 | 33 | 8.57E-06 |

### RSRC1 and RBM11 directly bind *MAPT* pre-mRNA and alter *MAPT* splicing *in vitro*

In order to prioritize SFs/RBPs for validation, we identified genes known to be expressed in brain (queried through the GTex portal; www.gtexportal.org) and neurons (Barres RNA-seq browser; www.brainrnaseq.org), as we predicted these would be most relevant to the regulation of *MAPT*, which is primarily neuronally expressed. This resulted in a panel of seven exon 2 includers/exon 10 excluders (Table 1, bold text).

To validate the association of candidate SF/RBPs with *MAPT* splicing, we first assessed whether they were able to directly interact with *MAPT* pre-mRNA in the context of human brain tissue. We carried out RNA pull-downs using desthiobiotinylated *in vitro* reverse transcribed RNA generated from a mini-gene containing *MAPT* exons 9-11, with full intervening intronic sequences (LI9LI10)^42^ and probed protein lysates derived from postmortem human prefrontal cortical tissue. The resulting eluates were examined by western blot for SFs/RBPs of interest. LI9LI10-derived *MAPT* pre-mRNA pulled down significant proportions of RSRC1 (41.62%, *p* < 0.001) and RBM11 (11.11%, *p* < 0.01) proteins from human brain lysates (Figure 2A-B), as well as a minimal, but not significant proportion of THOC3 (5.77%) and SNRNP25 (4.77%), suggesting that these factors may regulate *MAPT* exon 10 splicing via a direct interaction with its pre-mRNA.

**Figure 2.**
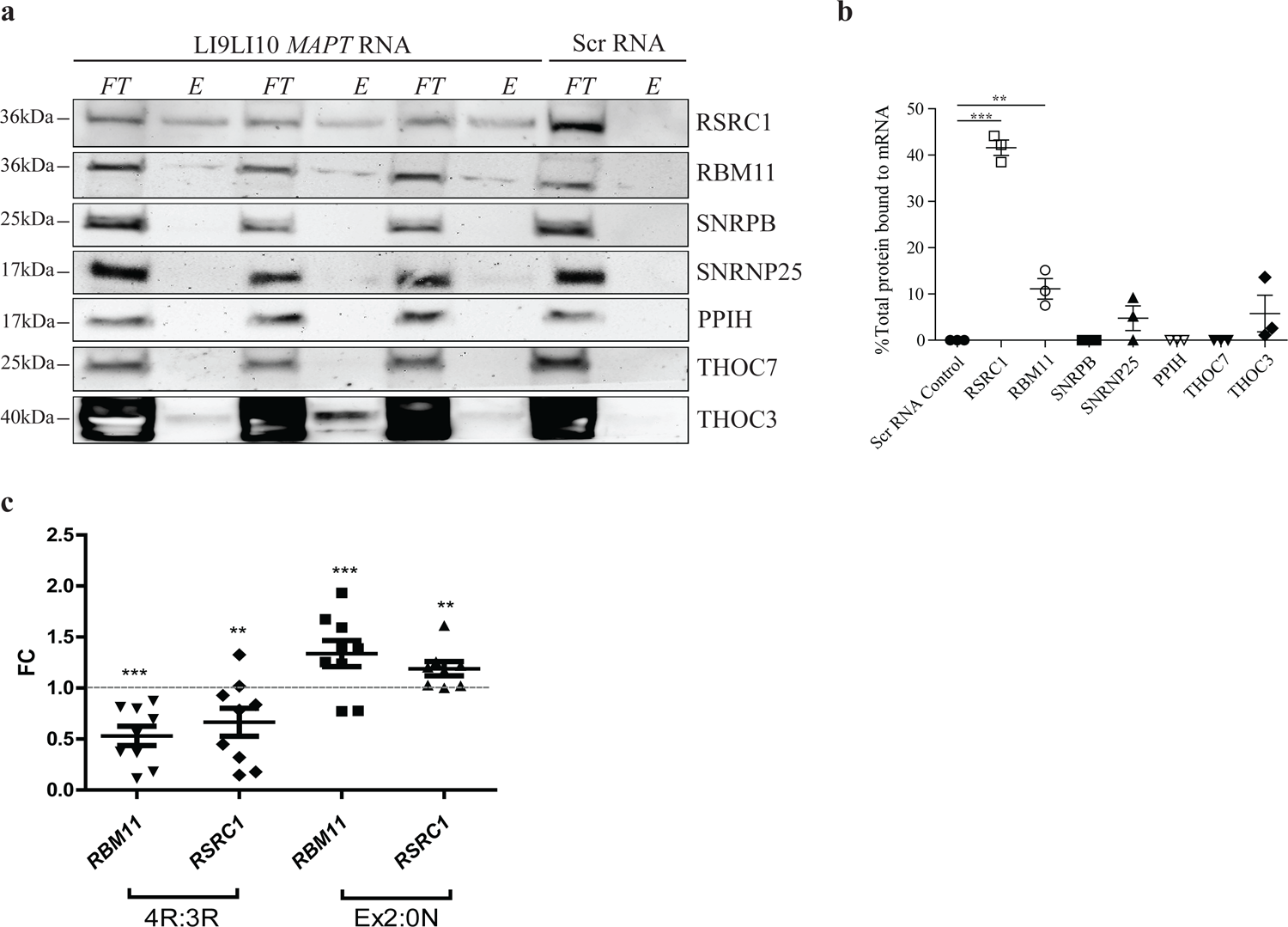
RBM11 and RSRC1 directly bind to *MAPT* pre-mRNA and regulate splicing **A)** *MAPT* pre-mRNA pull-downs from human brain tissue protein lysates for each target SF/RBP using the LI9LI10 minigene sequence or scrambled (Scr) RNA sequence as bait. *FT* = flow through fraction not bound to *MAPT* pre-mRNA or non-specific control, *E* = eluate fraction bound to target RNA. **B)** Quantification of RNA pull-down western blots in ***A***. Fraction of protein pulled down in *E* normalized to total protein present in *FT* and *E* fractions combined. N=3, each SF/RBP compared to scrambled control using t-test. \*\**p* < 0.01, \*\*\**p* < 0.001, ns = not significant. **C)** Fold change (FC) expression of 4R:3R and Ex2:0N ratios in SH-SY5Y cells by qRTPCR following overexpression of either *RBM11* or *RSRC1.* Asterisks denote significantly different expression compared to empty vector control, represented by grey line intersecting plot at FC = 1. One-way ANOVA and post-hoc Bonferroni tests for multiple comparisons. N = 3 independent experiments with 3 replicates each. \*\**p* < 0.01, \*\*\**p* < 0.001.

To functionally validate whether *RSRC1* and *RBM11* could alter *MAPT* splicing *in vitro*, we overexpressed these SF/RBPs in human neuroblastoma SH-SY5Y cells as an immortalized cell line proxy to neuronal cells (Figure S3A-B). As expression of *MAPT* exon 10 was very low in this cell line, we co-expressed the LI9L10 minigene with each SF/RBP to facilitate measurement of exon 10 exclusion. Consistent with their putative roles as exon 10 excluders, overexpression of either *RBM11* or *RSRC1* significantly reduced the 4R:3R ratio, as measured by qRTPCR (*RBM11 p* < 0.001, *RSRC1 p* < 0.01) (Figure 2C). While *PPIH* and *SNRPB* did not directly bind *MAPT* pre-mRNA, their overexpression resulted in a significantly reduced 4R:3R ratio in SH-SY5Y cells (*SNRPB p* < 0.01, *PPIH p* < 0.001) (Figure S3C). There was no significant effect of *THOC7*, *THOC3* or *SNRNP25* (Figure S3C), possibly due to poor overexpression efficiency. We then measured the ability for candidate SF/RBPs to alter N-terminal splicing of endogenous *MAPT*, and found that both *RBM11* and *RSRC1* overexpression significantly increased the Exon 2/0N ratio in SH-SY5Y cells (*RBM11 p* < 0.001, *RSRC1 p* < 0.01) (Figure 2C), consistent with their hypothesized role as exon 2 includers. In contrast, we did not see any effect of other candidate SF/RBPs on exon 2 inclusion (Figure S3D). We therefore conclude that *RBM11* and *RSRC1* may be important regulators of combinatorial N- and C-terminal *MAPT* splicing.

### Regulators of *MAPT* N-terminal splicing and *MAPT* exon 2 are differentially expressed in PSP and AD brain

Altered *MAPT* C-terminal splicing is a characteristic of several tauopathies^18^, including PSP^19^ and AD, for which there is inconsistent data^11, 13, 24–27^. We therefore queried the AMP-AD MAYO temporal cortex RNA-seq dataset, which includes both AD and PSP cases, in order to investigate whether *MAPT* splicing regulation may be altered in tauopathy brain. We calculated the fold change (FC) expression of every significant SF/RBP from our initial correlational analysis (Table S2) in PSP and AD compared to controls (Figure 3A). While PSP and AD shared largely similar patterns of SF/RBP expression dysregulation compared to controls, there were many genes that exhibited differential patterns of expression in either disease (Figure 3A). The group of SFs/RBPs that exhibited increased expression in PSP and reduced expression in AD comprised 4/7 of our candidate SF/RBPs, including both *RBM11* and *RSRC1,* which were associated with both exon 2 inclusion and exon 10 exclusion. In contrast, SFs/RBPs that were increased in AD and reduced in expression in PSP were significantly enriched for exon 2 excluders (Fisher’s exact test *p* = 0.0005) (Figure 3A). Furthermore, the net fold change (FC) expression of all exon 2 includers was significantly higher in PSP temporal cortex compared to AD (PSP average FC = 0.024, AD average FC = −0.036, *p* < 0.01) (Figure 3B), suggesting differential regulation of *MAPT* exon 2 splicing between diseases. To confirm this hypothesis, we compared *MAPT* exon 2, 3 and 10 PSI values in control, PSP and AD brain from the same dataset (Figure 3D-F). Consistent with the observed patterns of exon 2 splicing regulator expression in PSP and AD, we observed significantly different exon 2 inclusion across cases compared to controls (F_(2,134)_ = 4.01, *p* = 0.02), with significantly increased exon 2 PSI in PSP brain compared to controls (Tukey HSD *p* = 0.02), and a trend towards reduced exon 2 PSI in AD brain compared to controls (Figure 3D). In contrast, there was no significant difference in exon 3 or exon 10 PSI values between either disease and controls (Figure 3E-F). Taken together, this suggests there is differential dysregulation of *MAPT* N-terminal splicing regulators between AD and PSP brain, which results in altered expression of *MAPT* exon 2.

**Figure 3.**
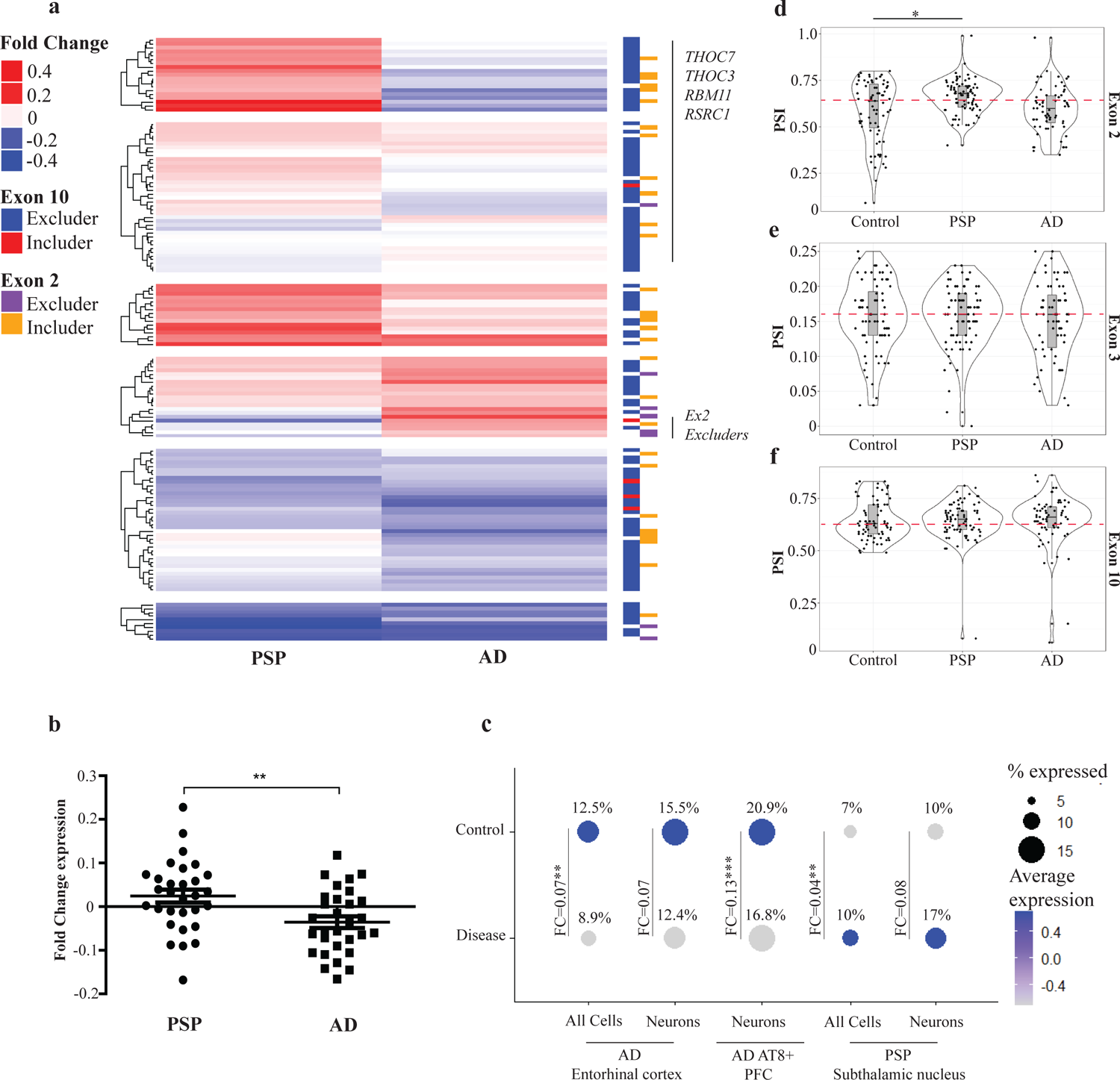
*MAPT* exon 2 expression and regulation is differentially altered in PSP and AD brain **A)** Fold change expression of each significant SF/RBP from the analysis in Figure 1G in PSP and AD brain compared to controls (AMP-AD MAYO temporal cortex), with unsupervised hierarchical clustering. Red indicates increased expression compared to controls, blue indicates reduced expression compared to controls. The direction of association of each SF/RBP with *MAPT* exon 2 and exon 10 splicing is indicated to the right of the figure. Clusters containing target SF/RBPs of interest from prior analyses and *MAPT* exon 2 excluders (purple) are indicated. **B)** Sum of the fold change expression of all exon 2 includers in PSP and AD brain compared to controls (AMP-AD MAYO temporal cortex). Statistical comparison between sum expression in PSP compared to AD brain, t-test, Error bars ±SEM. \*\**p* < 0.01. **C)** Single nuclei and single soma *RSRC1* expression in AD entorhinal cortex, AT8 positive/negative neurons and PSP subthalamic nucleus. FC = fold change expression disease/AT8 positive compared to controls/AT8 negative. Depth of color indicates scaled average expression, size of dot indicates proportion of cells expressing *RSRC1*, also denoted by percentage value above each dot. MAST linear model with Bonferroni correction, \*\**p* < 0.01, \*\*\**p* < 0.001 **D-F)** PSI values for exons 2 (***D***), 3 (***E***) and 10 (***F***) in control, AD and PSP brain (AMP-AD MAYO temporal cortex). Red dashed line indicates mean PSI value in controls for each exon. One-way ANOVA with post-hoc Tukey tests. Error bars ±SEM. \**p* < 0.05.

### *RSRC1* is differentially expressed in AD and PSP neurons in disease-relevant brain regions

While MAYO temporal cortex bulk RNA-seq data indicated altered expression of exon 2 splicing regulators in PSP and AD brain, these data may be influenced by altered cell type proportions in the disease context. Therefore, in order to validate neuronal and disease-specific patterns of expression of exon 2 splicing regulators, we assessed single nuclei sequencing (snuc-seq) from AD entorhinal cortex^43^ and PSP subthalamic nucleus^44^, as well as single-soma sequencing of hyperphosphorylated tau (AT8) positive neurons from AD prefrontal cortex^45^. We found that while *RBM11* was not detectable in AD snuc-seq data, and was expressed at very low levels in the other datasets, *RSRC1* was consistently detected and more highly expressed across data sets. While *RSRC1* expression was highest in microglia in both entorhinal cortex and subthalamic nucleus, we confirmed that it was also expressed in neurons in these regions (Figure S4A-B). Consistent with data from bulk temporal cortex tissue, we observed significantly lower *RSRC1* expression in AD tissue compared to controls (FC = 0.07, *p* < 0.01), as well as a lower proportion of *RSRC1*-expressing cells in AD compared to controls (8.9% vs 12.5%, respectively) (Figure 3C). We observed the same fold change expression and reduction in *RSRC1*-expressing cells when assessing neuronal populations specifically (Figure 3C), although due to the small proportion of neurons present in the data (Figure S4C), this was not significant. Interestingly, *RSRC1* expression was also higher in neurons derived from AD prefrontal cortex that were negative for hyperphosphorylated tau (AT8-), compared to AT8+ neurons (FC = 0.13, *p* = 3.33×10^-33^) (Figure 3C, Figure S4D), indicating that there may be an interaction between *RSRC1* expression, *MAPT* splicing regulation and the formation of tau pathology.

In contrast, we found significantly higher *RSRC1* expression in PSP subthalamic nucleus cells compared to controls (FC = 0.04, *p* < 0.01), as well as a higher proportion of *RSRC1*-expressing cells in PSP (10% compared to 7% controls) (Figure 3C). These differences were exacerbated when assessing neurons alone (FC = 0.08, 17% PSP neurons vs 10% control neurons) (Figure 3C), but similar to the AD data, the small proportion of neurons in this dataset (Figure S4E) precluded these data from reaching statistical significance. Interestingly, we were able to detect the opposite pattern of effect for the exon 2 excluders *QKI* and *PRPF38B* in AD and PSP neurons by snuc-seq: consistent with the temporal cortex bulk data, expression of *QKI* and *PRPF38B* were higher in AD neurons (*QKI* FC = 0.239, *PRPF38B* FC = 0.04) and lower in PSP neurons compared to controls (*QKI* FC = −0.03, 47.8% PSP vs 44.4% control neurons, *PRPF38B* FC = −0.125, 15.7% PSP vs 12.5% control) (Figure S4F). These data therefore support our assertion of differential expression of *MAPT* exon 2 regulators between AD and PSP brain. Furthermore, this demonstrates that SF/RBP expression is altered in neurons in disease-relevant brain regions.

### *MAPT* N-terminal isoforms are expressed at different ratios in AD and PSP brain

As we found evidence of coordinated splicing between *MAPT* exons 2 and 10, but observed differential expression of only exon 2 and exon 2 regulators in AD and PSP brain, we hypothesized that there may be loss of coordinated combinatorial splicing regulation between N- and C-terminal *MAPT* in tauopathy brain. We therefore carried out targeted *MAPT* isoform (iso)-seq on temporal cortex tissue from control, AD and PSP cases to assess the expression of full length transcripts (Figure 4A-C). We observed similar ratios of expression for each isoform as previously described by western blot analyses^12^: 1N3R and 0N3R were the two most highly expressed isoforms (∼30% and 25%, respectively), followed by 0N4R (∼21%) 1N4R (∼15%) and finally very low expression of both 2N isoforms 2N3R (∼1.4%) and 2N4R (∼0.8%) (Figure 4A). This pattern of expression was largely similar in AD and PSP cases, however there was a trend towards increased expression of both 0N isoforms in AD brain compared to controls (AD 0N3R 42.5% ± 12.7% vs 25% ± 5% controls, AD 0N4R 31.6% ± 3% vs 21% ± 4.6% controls), although due to high variability, these differences did not reach statistical significance (Figure 4A). We then compared the 4R:3R ratios for each N-terminal isoform, and found that despite increased expression of 0N isoforms in AD, there was no difference for either AD or PSP in the 0N4R:0N3R ratio (Figure 4B). In contrast, we observed a trend towards an increase in exon 2-containing 4R:3R ratios in PSP cases compared to controls (1N4R:1N3R PSP = 1.1 ± 0.38 vs 0.53 ± 0.07 controls, 2N4R:2N3R PSP = 0.8 ± 0.13 vs 0.55 ± 4.9 controls) (Figure 4B), consistent with both our observation of increased exon 2 inclusion in temporal cortex (Figure 3D) and the known pathological accumulation of 4R tau isoforms in PSP pathology. Lastly, when comparing N-terminal isoform expression, we found no differences in PSP brain, but a trend towards reduced 1N/0N and 2N/0N ratios in AD brain specifically (1N/0N AD = 0.43 ± 0.13 vs 1.2 ± 0.33 controls, 2N/0N AD = 0.02 ± 0.01 vs 0.05 ± 0.01 controls) (Figure 4C), likely due to the increase in expression of 0N isoforms (Figure 4A). Interestingly, we also observe an increase in the 2N/1N ratio in AD brain (0.1 ± 0.03 vs 0.05 ± 0.005 controls) (Figure 4C), which may be due to slightly decreased 1N expression or increased 2N3R expression we observe in AD compared to controls (Figure 4A).

**Figure 4.**
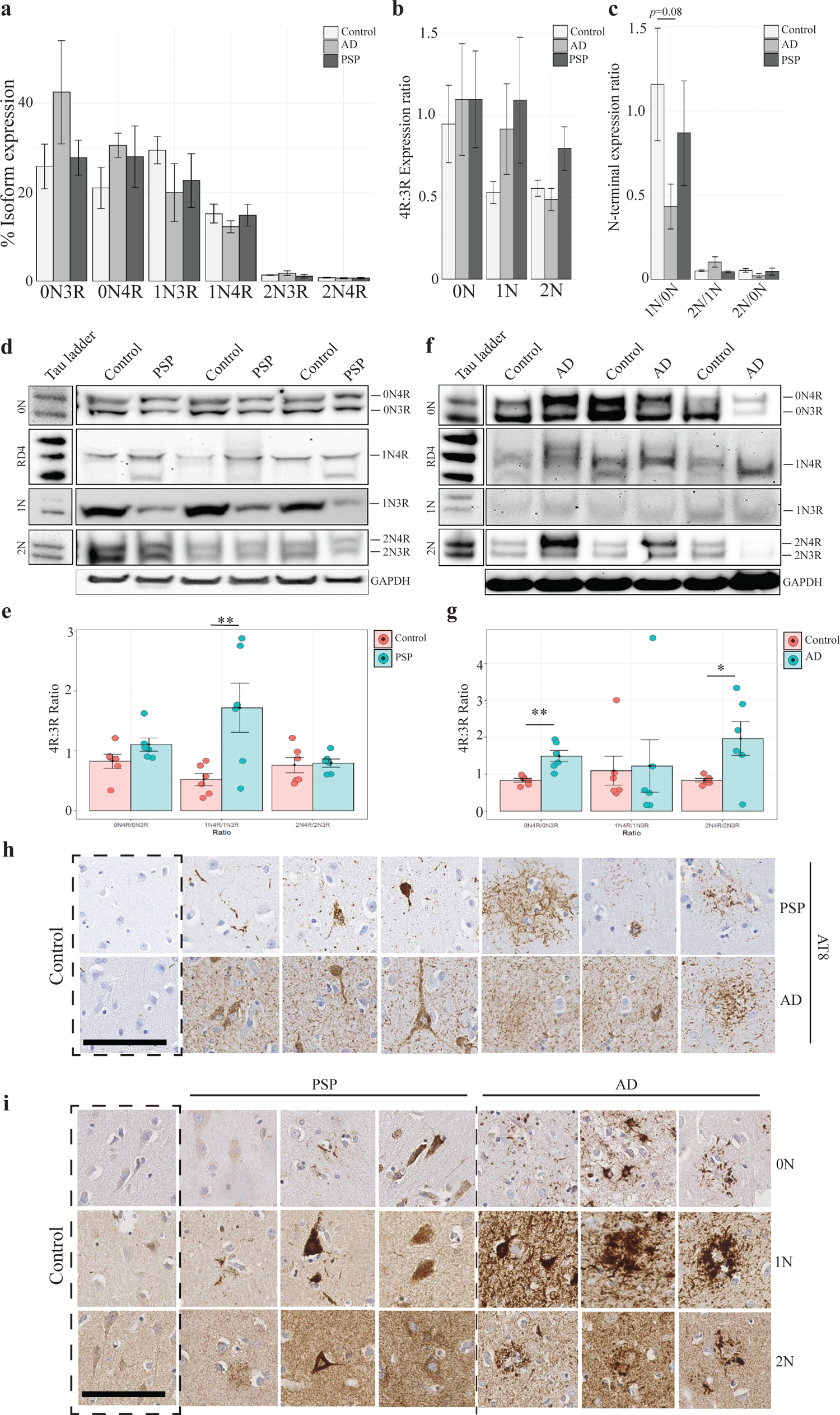
Coordination of N- and C-terminal splicing regulation is altered in AD and PSP brain, and is associated with tau pathology. **A-C)** Proportion of each full length *MAPT* isoform (***A***), 4R:3R ratio for each N-terminal isoform (***B***) and N-terminal isoform ratios (***C***) as detected by targeted *MAPT* iso-seq in human control, AD and PSP temporal cortex. One-way ANOVA with post-hoc Tukey tests, all comparisons did not reach statistical significance. Error bars ±SEM. **D)** Dephosphorylated tau isoform expression in PSP temporal cortex compared to tau ladder detected by specific N-terminal antibodies against 0N, 1N and 2N. Anti-1N tau showed poor detection for 1N4R isoforms, so anti-4R tau antibody RD4 was used for detection of this isoform. GAPDH used as loading control. **E)** Quantification and 4R:3R ratio of N-terminal isoforms in ***D***. Each point denotes a different individual brain lysate, N = 6. Error bars ±SEM. Student’s t-test \*\**p* < 0.01 **F)** Dephosphorylated tau isoform expression in AD temporal cortex compared to tau ladder with same antibody detection as in ***D***. GAPDH used as loading control. **G)** Quantification and 4R:3R ratio of N-terminal isoforms in ***F***. Each point denotes a different individual brain lysate, N = 6. Error bars ±SEM. Student’s t-test \**p* < 0.05, \*\**p* < 0.01 **H)** Representative images of labeling of control, AD and PSP temporal cortex with marker of hyperphosphorylated tau, AT8, indicating different tau pathologies present across cases. Sections counterstained with hematoxylin and eosin. N=3-4. Scale bar = 100µm. **I)** Representative images of tau pathology labeling in control, AD and PSP temporal cortex with anti-0N, 1N and 2N tau antibodies from several individuals. Sections counterstained with hematoxylin and eosin. N=3-4. Scale bar = 100µm.

To examine whether these transcriptional changes were apparent at the protein level, we carried out western blot analyses in the same tissues from the same individuals (Figure 4D-G). Consistent with the iso-seq data, there was a significantly increased 1N4R:1N3R ratio in PSP brain compared to controls (control 1N4R:1N3R ratio = 0.52 (SEM= 0.1), PSP 1N4R:1N3R ratio = 1.7 (SEM = 0.4), *p* = 0.009), which was largely driven by a reduction in 1N3R tau (Figure 4D-E). In contrast, there was no difference in either 0N or 2N 4R:3R ratios in PSP (Figure 4D-E). In AD brain, there was an accumulation of soluble 0N and 2N isoforms by western blot, consistent with the iso-seq data (Figure 4C, F-G). When examining 4R:3R ratios for each N-terminal isoform, we observed significantly increased 0N4R:0N3R (control ratio = 0.83 (SEM = 0.05), AD ratio = 1.49 (SEM = 0.15), *p* = 0.002) and 2N4R:2N3R ratios (control ratio = 0.83 (SEM = 0.05), AD ratio = 1.96 (SEM = 0.46), *p* = 0.03) in AD brain compared to controls (Figure 4F-G), indicating that while there were no transcriptional differences in the 4R:3R ratio for these isoforms, there may be impaired degradation of 0N4R and 2N4R tau in AD resulting from their increased expression.

### Pathologically aggregated tau in PSP and AD brain are associated with different N-terminal isoforms

While western blot analysis of AD and PSP temporal cortex revealed differences in the accumulation of soluble N-terminal tau isoforms, we wanted to determine whether the formation of different neuropathological features between tauopathies may be due to the insoluble aggregation of different N-terminal tau isoforms. After validating the specificity of each N-terminal antibody by overexpressing different tau isoforms in N2a cells (Figure S5A-B), we carried out immunohistochemistry (IHC) on control, AD and PSP brain sections from the temporal cortex for hyperphosphorylated pathogenic tau (AT8; Figure 4H)) and each N-terminal isoform (Figure 4I). We observed the anticipated hyperphosphorylated tau neuropathology in both PSP and AD brain (Figure 4H), specifically the widespread presence of neurofibrillary tangles and neuropil threads throughout the AD cortex, and sparse glial plaques and neurofibrillary tangles in PSP tissue. In comparison, there was little to no signal in control brain (Figure 4H).

There was little signal for 0N tau in PSP brain (Figure 4I), although in one case we observed sparse labeling of neuropil threads or possible glial involvement (Figure 4I). In AD brain the 0N antibody did not label neurofibrillary tangles, but we did observe labeling of neuropil threads and dystrophic neurites surrounding amyloid plaques (Figure 4I). Interestingly, in one case the 0N antibody labelled thorny astrocytes consistent with an age-related tau astrogliopathy (ARTAG) pathology that was not visible with the AT8 antibody (Figure 4H-I). While it is surprising that there was little immunolabeling of 0N Tau in human brain, given the high levels of 0N transcripts, this pattern was replicated with a second antibody against 0N Tau (Figure S5C).

1N Tau immunostaining was primarily present in neurofibrillary tangles and pre-tangles (Figure 4I) throughout the cortex in all three PSP cases, as well as neuropil granules and threads. In AD, this isoform was also the most prevalent in neurofibrillary tangles (Figure 4I), but was primarily present in dystrophic neurites surrounding amyloid plaques (Figure 4I). Finally, while 2N Tau was observed in some neurofibrillary tangles in PSP and AD brain (Figure 4I), these were less common than the 1N labelled neurons. 2N Tau was also observed in dystrophic neurites surrounding amyloid plaques in AD brain, but this labelling was far less dense than the 1N Tau and less tightly co-localized with plaques (Figure 4I). A second 2N antibody revealed a similar labeling of neurofibrillary tangles in AD and PSP brain, but less involvement in dystrophic neurites (Figure S5C).

In order to directly compare the accumulation of different tau N-terminal isoforms in pathogenic inclusions, we carried out Opal multiplex labelling of adjacent brain sections from the same individuals using all three N-terminal antibodies in conjunction with AT8 and β-amyloid staining (Figure 5A-B, Figure S6A-B). Consistent with the IHC staining, we observed primarily 1N and 2N tau in AT8-positive neurofibrillary tangles in PSP temporal cortex, but all three N-terminal isoforms were present in AD-associated tangles (Figure 5A-B, Figure 6A-B). Dystrophic neurites surrounding amyloid plaques were primarily associated with 0N and 1N accumulation, with less 2N involvement (Figure 5A, Figure S6A). In contrast, amyloid plaques present in either controls or PSP cases were not associated with any tau staining (Figure S6B). Interestingly, 2N tau was absent in glial pathology observed in both AD and PSP cases, suggesting that 2N tau is unable to accumulate in glia, while astrocytic tufts in PSP brain were labelled primarily by 1N tau (Figure 4I), consistent with the increased accumulation of 1N4R tau isoforms we observed by western blot and transcriptomic analyses.

**Figure 5.**
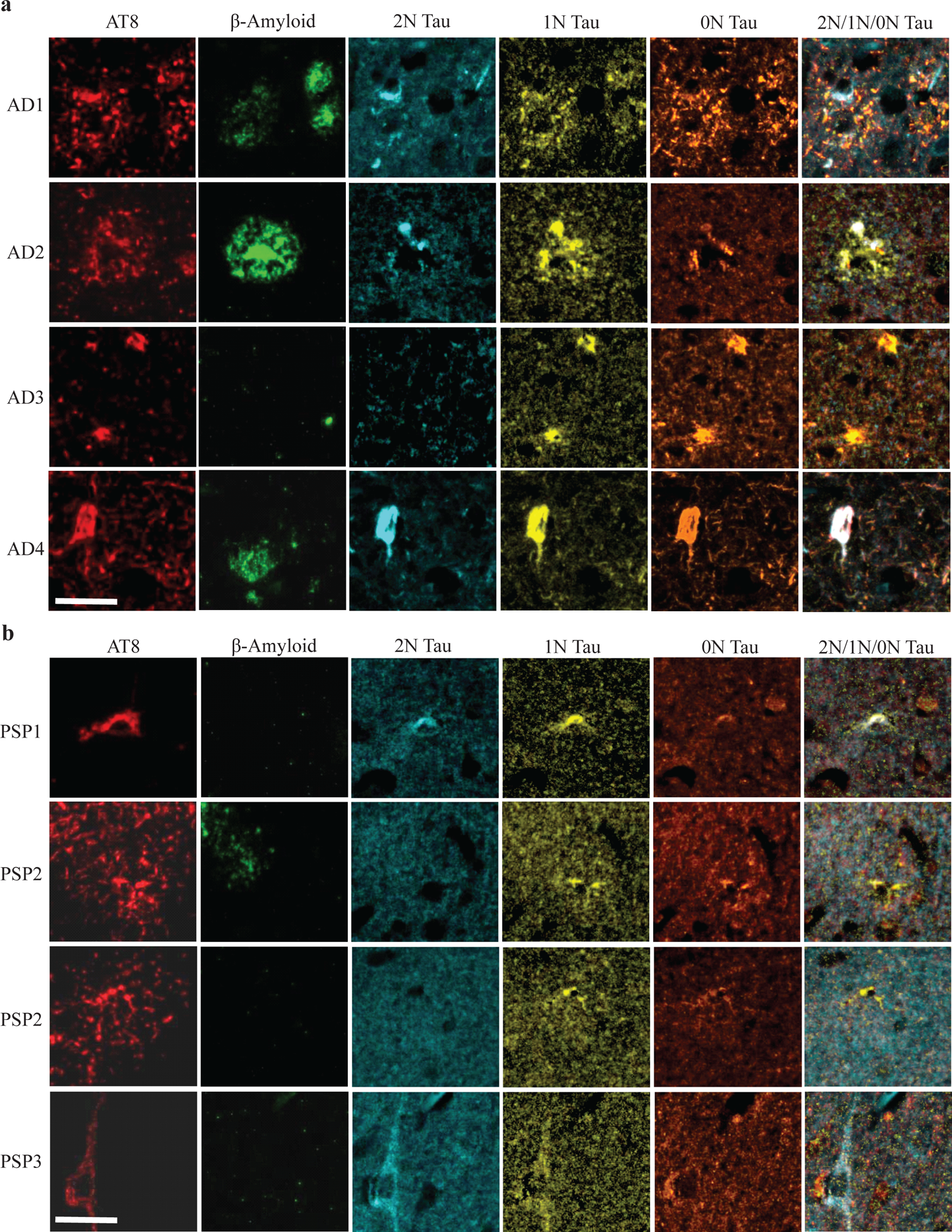
N-terminal isoforms accumulate differently in AD and PSP neuronal and glial pathologies **A)** Representative images of multiplex immunofluorescent labeling of AD temporal cortex with AT8 (red), β-amyloid (green), 2N tau (blue), 1N tau (yellow) and 0N tau (orange), and overlay of all three N-terminal tau antibodies in 4 different individuals. Examples of tau accumulation in dystrophic neurites can be found in AD1 and AD2, thorny astrocytes present in AD3, and an example of a neurofibrillary tangle shown in AD4. N = 4, scale bar = 50µm. **B)** Representative images of multiplex immunofluorescent labeling of PSP temporal cortex from 3 individuals as in ***A***. Examples of early and pre-tangles shown in PSP1 and PSP3, while examples of astrocytic tufts are shown in PSP2. N = 3, scale bar = 50µm.

## DISCUSSION

To date, the contribution of N-terminal *MAPT* splicing to disease pathogenesis has largely been overlooked compared to evaluation of exon 10 splicing and 4R tau expression. However, N-terminal tau is relevant to disease pathogenesis: for example, N-terminal fragments are prominent in AD cerebrospinal fluid (CSF), and their secretion from cells can be inhibited by the presence of exon 2^15, 46^. In contrast, 0N Tau is more readily cleaved and released from human neuroblastoma cells^46^. This is consistent with our IHC data, where we observe little 0N tau accumulation in neurons, but prominent 1N and 2N tau in neurofibrillary tangles. The regulation of tau release by N-terminal inserts also has implications for our understanding of tau spread and seeding, as different isoforms may be available extracellularly, and exhibit different seeding competencies. Furthermore, aberrant folding of N-terminal tau is one of the earliest pathological changes identified in tauopathies^47^, thus supporting the assertion that N-terminal splicing is a relevant consideration when modeling and investigating tauopathy.

N-terminal tau splicing may play a role in modifying tau subcellular localization and aggregation propensity. The N-terminal contains a plasma-interacting domain^48^ that interacts with synaptic proteins and Annexin A6^49, 50^, which results in retention of tau in the axonal compartment. While exons 2 and 3 are not within this domain, it is possible that they modify these interactions and impact the subcellular localization of tau, resulting in the increased propensity for certain isoforms to accumulate in these regions. Indeed, murine 0N tau has been found to localize in axons, whereas 1N was enriched in dendrites and 2N tau was depleted from cytoskeletal structures^51^, indicating that N-terminal splicing likely modifies tau function. 1N tau has also been found to more readily accumulate and aggregate: the presence of exon 2 promotes the fibrillary extension of tau filaments *in vitro*^52^ and exon 2-containing proteins more readily polymerize than 0N tau^53^, therefore increased exon 2 inclusion would be likely to worsen tau aggregation. This is consistent with our observation of strong 1N tau immunostaining in PSP astroglia and tangles in both PSP and AD neurons. Despite this, we observe increased 0N expression in AD brain. However, the resulting protein accumulation was soluble, and associated with dystrophic neurites surrounding amyloid plaques rather than in neurofibrillary tangles. N-terminal tau splicing has also been found to be required for specific interactions with proteins associated with synaptic signaling and the plasma membrane^54^, supporting the assertion that different N-terminal isoforms likely facilitate distinct cellular functions. Characterizing the disruption of N-terminal splicing is therefore important for understanding tau biology and the mechanisms underlying disease pathogenesis.

We have identified coordinated regulation of *MAPT* exons 2 and 10, indicating that N- and C-terminal splicing of either region likely does not occur fully independently of the other. This phenomenon has been reported once previously using polony-based exon profiling^13^. Disrupting this coordination will therefore lead to imbalanced expression of different tau isoforms. Interestingly, while the expression of numerous exon 10 splicing regulators were altered between AD, PSP and control brains, we observed enrichment of exon 2 regulators with opposing patterns of differential expression between PSP and AD. As anticipated from this pattern of expression, we observed increased exon 2 inclusion in PSP and reduced exon 2 in AD. However, assessment of full length isoforms by iso-seq and western blot revealed that shifts in N-terminal splicing were coupled to alterations in the 4R:3R ratio, supporting our hypothesis that loss of coordinated regulation between N- and C-terminal splicing contributes to tauopathy pathogenesis. The change in the 4R:3R ratio for specific N-terminal isoforms may explain why we did not observe an increase in exon 10 inclusion in PSP by short-read bulk RNA-seq; the most highly expressed 0N isoforms did not exhibit a change in the 4R:3R ratio in PSP brain, therefore the lack of change in 0N4R may have diluted out increases in 1N or 2N4R. Curiously, the increased 1N4R:1N3R ratio observed in PSP appeared to be largely due to a loss of 1N3R rather than an increase in 1N4R tau, thus raising the possibility that in combination to 1N4R tau accumulation, losing 1N3R tau expression and function could also be detrimental to neuronal health.

Aberrant regulation of splicing is a known phenotype of some forms of inherited tauopathy^55^, which is hypothesized to be due to the sequestration and mislocalization of SF/RBPs to the cytosol and into stress granules^55^. SF/RBP dysfunction is therefore a likely mechanism underlying aberrant *MAPT* splicing in non-familial AD and PSP. While we do not characterize the mechanism of SF/RBP disruption in AD and PSP brain here, we do identify differential expression of numerous SF/RBPs between AD and PSP compared to controls, which is suggestive of separate downstream effects of splicing dysregulation that ultimately contribute to the pathogenesis of either disease.

We found that *MAPT* splicing is likely regulated by numerous splicing factors, of which *RSRC1* and *RBM11* may be of particular interest. However, it is likely that there are many other modifiers that were not identified in our screen. Several of our SF/RBPs of interest, including *SNRPB*, *SNRNP25*, *THOC7* and *THOC3*, are known to be components of protein complexes that regulate splicing, and therefore would be unlikely to be isolated by the RNA pull-down assay. Indeed, overexpression of some of these SF/RBPs was able to significantly shift the 4R:3R ratio in human neuroblastoma cell lines. While expressed in brain and in neurons, many of these SF/RBPs are ubiquitously expressed in many different tissues and cell types. Indeed, we observed the highest expression of *RSRC1* in microglia in human brain snuc-seq data. Therefore, while their dysregulation may be impacting *MAPT* splicing in neurons, there will likely be wider effects of altered expression and splicing in other neural cell types that may also be relevant for disease pathogenesis.

*RSRC1* is a member of the serine and arginine rich-related protein family, which are highly conserved regulators of alternative splicing, but how *RSRC1* regulates this process is unknown. It is hypothesized that RSRC1 plays a role in 3’ splice site selection by interaction with splicing regulators U2AF and SRSF2^56^, although it also possible that it may interact directly with pre-mRNA via its RS (arginine-serine) domain. However, there is no specific RNA motif that has been characterized as binding RS domains, thus the region in which RSRC1 may be binding *MAPT* pre-mRNA is currently unknown. Mutations within this gene have been associated with intellectual disability^57, 58^, whereas the silencing of *RSRC1* in SH-SY5Y cells has been associated with downregulation of genes associated with schizophrenia, Alzheimer’s disease and dementia^58^, thus indicating the relevance of this gene for brain function.

*RBM11* is a brain specific splicing factor that exhibits fluctuating expression with brain development, with high expression throughout embryogenesis, peaking at perinatal days 0-3^59^, after which *MAPT* exon 10 expression increases^8^, consistent with our observations that *RBM11* expression may promote exon 10 exclusion. *RBM11* is associated with the choice of 5’ splice sites and may antagonize the activity of other SF/RBPs such as *SRSF1*^59^, potentially regulating *MAPT* splicing by direct interaction with pre-mRNA and by inhibiting binding of competing SFs/RBPs. *RBM11* is downregulated in the PS19 mouse model of tauopathy^55^, consistent with the direction of effect we observe in AD brain. However, this mouse model expresses only 1N4R tau and as such the effect of *RBM11* disruption on *MAPT* splicing was not measurable in this model. It should be noted that we were unable to validate *RBM11* expression in snuc-seq data as its expression was very low, therefore the relevance it may have to tauopathy and *MAPT* splicing requires additional investigation.

We were unable to identify regulators of *MAPT* exon 3 splicing from these data, or to compare its regulation between 17q21.31 H1 and H2 haplotypes due to the low frequency of H2 carriers. Increased Exon 3 inclusion is consistently identified on the H2 background^12, 39^, an effect we also observe in these data. However, the mechanism underlying increased exon 3 inclusion, and whether its expression is protective against tauopathy is currently unknown, although it may reduce the fibrillization of tau^52^. It should be noted that exon 3, and subsequently 2N tau expression, is very low in adult human brain (accounting for less than 3% of all transcripts in our targeted iso-seq data), therefore the extent to which it may contribute to disease is unclear. Nevertheless, we observe some alterations in 2N expression and accumulation in AD and PSP, the most striking of which was the absence of 2N tau in glial pathology, indicating that 2N tau is either not released from neurons or is not internalized or aggregated by astrocytes. These data have implications for the design of experimental models of tauopathy where a single tau isoform is expressed in order to ensure the most disease and pathology appropriate isoform is utilized.

In conclusion, we propose that changes in SF/RBP expression result in differential splicing of the *MAPT* N-terminus between AD and PSP, resulting in the expression of isoforms with different aggregation properties and subcellular localizations, thus explaining the distinct neuropathological phenotypes of each disease. It would therefore be of great interest to investigate the role of N-terminal splicing in other primary tauopathies associated with different pathologies, such as Pick’s disease (PiD), primary age-related tauopathy (PART), age-related tau astrogliopathy (ARTAG) and chronic traumatic encephalopathy (CTE) to determine whether these diverse disorders also exhibit loss of *MAPT* exon 2 and 10 splicing coordination. These data indicate that it is unlikely that exon 10 splicing is alone in underlying and regulating disease pathogenesis and tau neuropathology in either AD or PSP, but rather the combinatorial expression of specific and N- and C-terminal *MAPT* isoforms is relevant for understanding the development of tauopathy. While alterations in the 4R:3R ratio are undoubtedly important and relevant to our understanding of tau pathogenesis, N-terminal splicing is likely to be an important modifier of disease and pathology, and should be carefully assessed.

## METHODS

### RNA-seq data analysis

Aligned BAM files and gene expression count data were downloaded from the AMP-AD consortium through Synapse (https://adknowledgeportal.synapse.org/). *MAPT* exon level counts were calculated using the FeatureCounts feature within the Subread package^60, 61^, and PSI values were determined using the Mixture of Isoforms (MISO) package^62^. For statistical analysis, associations between *MAPT* PSI values and the expression of SFs/RBPs were carried out using a linear model in R with RNA integrity number (RIN), postmortem interval (PMI), sex, and age at death included in the model as covariates. The resulting *p*-values were Bonferroni-corrected for the number of comparisons. For heatmap plotting, correlation coefficients were generated using Pearson’s correlation as part of the cor function in R. For the comparison of SF/RBP expression in PSP and AD brain, the fold change expression of each SF/RBP in disease brain was calculated in comparison to controls and plotted in R using ComplexHeatmap^63^, and statistically significant differences were determined by linear regression of expression values including the previously described covariates.

At the time of analysis, genotype data were unavailable for the MSBB cohort, so 17q21.31 haplotype was determined by Taqman genotyping. The relevant DNA was obtained from the NIH Neurobiobank. Taqman genotyping was carried out for H2 tag SNPs rs8070723 and rs1052553 using commercially available assays. Haplotypes were determined for the other cohorts using the same tag SNPs from genotype data downloaded from the AMP-AD knowledge portal (https://adknowledgeportal.synapse.org/).

### Single-nuclei and single-soma sequencing analysis

Snuc-seq processed gene counts and covariates derived from AD and control entorhinal cortex^43^ were downloaded from the Gene Expression Omnibus (GEO) (GSE138852). Data were further normalized and analyzed in Seurat 3.0 dev^64, 65^. Data from different individuals were integrated and scaled using SCTransform^66^ while regressing out the percentage of mitochondrial genes, the number of genes per cell and the number of reads per cell. Principal components analysis (PCA) was carried out in Seurat using the top 3000 most variable genes, and data was reduced using UMAP^67^. Cell types present within each cluster were already annotated in the downloaded metadata. Differential gene expression analysis was carried out between AD cases and controls across the whole data set, or within the neuronal cluster only, using the MAST model applied to log normalized raw count data, including the percent of mitochondrial genes as a covariate.

Aligned HDF5 feature barcode matrices for the single-soma sequencing data of AT8 positive and negative neurons from AD PFC^45^ were downloaded from GEO (GSE129308) and processed in Seurat^64, 65^. Data were filtered for cells expressing > 200 genes, < 2500 reads and < 10% mitochondrial genes. Data were integrated, transformed and reduced as described above. Differential gene expression analysis was carried out between AT8 positive and AT8 negative cells across the whole data set, using the MAST model applied to log normalized raw count data, including the percent of mitochondrial genes, age, RNA integrity number (RIN) and postmortem interval (PMI) as covariates.

Aligned HDF5 feature barcode matrices for PSP snuc-seq data^44^ were kindly shared by Drs. Pereira and Crary, and were filtered, integrated, transformed and reduced in the same manner as the AD snuc-seq and AT8 soma-seq data described above. Individual clusters were identified in Seurat using the default resolution factor 0.5. Cell types within each cluster were defined using visualization of specific markers utilized in the AD snuc-seq data^43^; *CD74* (microglia), *AQP4* (astrocytes), *MEGF11* (oligodendrocyte precursor cells (OPCs)), *MOBP* (oligodendrocytes), *SYT1* (neurons) and *FLT1* (endothelial cells). Differential gene expression analysis was carried out between PSP cases and controls across the whole data set, or within the neuronal cluster only, using the MAST model applied to log normalized raw count data, including the percent of mitochondrial genes and age as covariates.

### Human brain tissue

Fresh frozen human control, AD and PSP temporal cortices were acquired from the Mount Sinai Neuropathology Core brain bank and the Harvard Brain Tissue Resource Center, University of Maryland Brain and Tissue Bank and Mount Sinai Brain Bank via the NIH Neurobiobank. Formalin fixed paraffin embedded sections from temporal cortex were acquired from the Mount Sinai Neuropathology Core brain bank, with neuropathological diagnosis being determined by Dr. John Crary. All post-mortem tissues were collected in accordance with the relevant guidelines and regulations of the respective institutions. A summary of tissues used in this project are described in Table S3.

### Cell culture

All cell culture reagents were purchased from Thermo Fisher Scientific, unless otherwise stated. SH-SY5Y cells were grown in IMDM media supplemented with 1% penicillin/streptomycin and 10% FBS, and grown and maintained at 37°C with 5% CO_2_ in a humid environment. Prior to transfection, cells were seeded into 6 well plates at a density of 1.6×10^5^ cells per well. The next day, cells were transfected with 1.25ug of LI9LI10 mini-gene and 1.25ug of plasmid DNA of the SF/RBP of interest (all Origene) using Lipofectamine 3000. Cells were collected for RNA extraction and analysis 48 hours later. For N-terminal antibody validation, N2a cells were transfected with 2.5ug of either a 0N3R, 1N3R or 2N3R *MAPT* cDNA vector (Origene), and after 48 hours were either fixed with 10% formalin (Sigma Aldrich) for 15 minutes at room temperature for immunofluorescence, or pelleted in PBS for protein extraction and western blotting.

### RNA pull-down

The LI9LI10 mini-gene was digested by NotI (Cell Signaling Technologies) and SgfI (Promega) to excise the *MAPT* coding sequence and upstream T7 promoter from the PCI-Neo backbone. The resulting DNA was transcribed *in vitro* using the T7 Megascript kit (ThermoFisher Scientific), incubated at 37°C for 4 hrs, followed by 15 minutes treatment with DNase to degrade any remaining DNA template. The resulting RNA was isolated by Lithium Chloride precipitation, and examined on a 1% agarose gel for the anticipated product size, compared against Lambda DNA digested with HindIII and EcoRI (both Cell Signaling Technologies). RNA was labelled using the 3’ end desthiobiotinylation RNA labelling kit (ThermoFisher), with 150ug/25nM RNA per reaction incubated at 16°C overnight. Labelled RNA was then isolated by chloroform and ethanol precipitation. A negative control consisting of a scrambled RNA sequence was labelled at the same time. Labelling efficiency was measured by comparison of chemiluminescent signal from labelled sample RNA with a positive control using the ThermoFisher Scientific Chemiluminescent Nucleic Acid detection module. 25nM of labelled RNA was then bound to nucleic acid-compatible streptavidin magnetic beads using the Pierce Magnetic RNA-Protein Pull-down kit (ThermoFisher Scientific) and incubated with 100ug protein lysate from human brain overnight at 4°C. Bound protein was eluted from the beads and immediately subject to western blot analysis for SFs/RBPs of interest.

### SDS-PAGE gel electrophoresis and western blot

Soluble protein was collected from cell pellets and human brain tissue by resuspension in Cell Lysis Buffer (Cell Signaling Technologies) supplemented with 10µM PMSF on ice. Cells or tissue were then sonicated briefly on ice and spun at 13,000xg for 10 minutes at 4°C to pellet debris. Protein concentration was determined by BCA assay (ThermoFisher Scientific). Prior to western blotting, human brain protein lysates were dephosphorylated in order to accurately determine Tau isoforms by size. Lysates were incubated with 100 units of Lambda protein phosphatase (LPP; Cell Signaling Technologies) per 10ug total protein, supplemented with 1x Protein MetalloPhosphatases buffer and 1mM MnCl_2_, and incubated at 30°C for 3 hours before being analyzed by western blot.

For SDS-PAGE gel electrophoresis, 10-30ug of protein was incubated with 1x reducing agent and 1x LDS sample buffer at 70°C for 10 minutes before immediately being loaded onto a BOLT 4-16% Bis-Tris gel (ThermoFisher Scientific) in 1x MES buffer. For splicing factor analyses, electrophoresis was carried out for 20 minutes at 200V before blotting. For tau isoform analyses, electrophoresis was carried out for 60 minutes at 100V before blotting. Gels were blotted onto nitrocellulose membranes using the iBlot system (ThermoFisher Scientific), and blocked for a minimum of 30 minutes in 5% milk in PBS-T. Primary antibodies were prepared at dilutions described in SI Table 4 in 5% milk in PBS-T and incubated with the membrane at 4°C overnight. Membranes were washed 3x in PBS-T, then incubated with either HRP Goat Anti-Rabbit or HRP Horse Anti-Mouse secondary antibodies (Vector laboratories) at a dilution of 1:20,000 in 5% milk in PBS-T for two hours at room temperature. Following three additional washes, membranes were then incubated with WesternBright ECL HRP substrate (Advansta) for 3 minutes before imaging on a UVP ChemiDoc. For re-staining, blots were incubated in Restore PLUS stripping buffer (ThermoFisher Scientific) for 15 minutes at room temperature, followed by one wash in PBS and re-blocking.

### qRTPCR

Cell pellets were collected by washing and scraping into ice-cold PBS. RNA was extracted from cell pellets using the Qiagen RNeasy mini RNA extraction kit, and reverse transcribed using the high capacity RNA-to-cDNA kit (ThermoFisher Scientific). qRTPCR for specific *MAPT* exons and isoforms was carried out using SybrGreen mastermix with the following primers: *MAPT* 4R Forward 5’-CGGGAAGGTGCAGATAATTAA-3’, Reverse 5’-GCCACCTCCTGGTTTATGATG-3’; *MAPT* 3R Forward 5’-AGGCGGGAAGGTGCAAATA-3’, Reverse 5’-GCCACCTCCTGGTTTATGATG-3’; *MAPT* 0N Forward 5’-TTTGAACCAGGATGGCTGAG-3’, Reverse 5’-ATGCCTGCTTCTTCAGCTTT-3’; *MAPT* Exon 2 Forward 5’-TTTGAACCAGGATGGCTGAG-3’, Reverse 5’-CTGCAGGGGAGATTCTTTCA-3’. SF/RBP overexpression and knockdown was validated and quantified by qRTPCR using commercially available Taqman assays (ThermoFisher Scientific).

### *MAPT* targeted iso-seq

RNA was extracted from human temporal cortex brain tissue as described above and submitted to the Icahn School of Medicine at Mount Sinai Genomics CoRE for single molecule real time (SMRT) isoform sequencing (iso-seq) on the PacBio RS II platform using the following primers: Forward 5’-ATG GAA GAT CAC GCT GGG AC-3’, Reverse 5’-GAG GCA GAC ACC TCG TCA G-3’. Raw sequencing reads were passed through the ISOseq3 pipeline to detect full-length transcripts expressed in each sample. Beginning with raw subreads, single consensus sequences were generated for each *MAPT* amplicon with a SMRT adapter on both ends of the molecule. SMRT adapter sequences were then removed and *MAPT*-specific primer sequences were identified to orient the isoforms. Isoforms were subsequently trimmed of poly(A) tails and concatemers were identified and removed. Isoform consensus sequences were then predicted using a hierarchical alignment and iterative cluster merging algorithm to align incomplete reads to longer sequences. Finally, clustered isoform sequences were polished using the arrow model and binned into groups of isoforms with predicted accuracy of either ≥ 0.99 (high quality) or < 0.99 (low quality). The resulting isoforms were aligned to hg38 using the GMAP aligner^68^ and isoform calling, collapsing and measurements of abundance were carried out using the Cupcake/ToFU pipeline (https://github/Magdoll/cDNA_Cupcake).

### Immunohistochemistry

Immunohistochemical staining was carried out on formalin fixed paraffin-embedded brain sections by the Neuropathology Brain Bank and Research CoRE at the Icahn School of Medicine at Mount Sinai using the Ventana BenchMark autostainer. Slides were scanned on a Leica SCN400 at 40x. A list of antibodies used and their relevant dilutions can be found in Table S4.

### OPAL multiplexed immunofluorescence

Multiplexed immunofluorescent staining was carried out using the Opal Polaris 7 color IHC detection kit (Akoya biosciences) according to manufacturer’s instructions. Briefly, slides were baked for 1 hour at 65°C, then deparaffinized with xylene and rehydrated with a graded series of ethanol concentrations. For epitope retrieval, slides were microwaved in AR buffer (provided with the OPAL IHC detection kit) for 45s at 100% power, followed by an additional 15 minutes at 20% power. After cooling, slides were blocked for 10 minutes in blocking buffer then incubated with the first primary antibody at room temperature for 30 minutes. Slides were rinsed three times in TBS-T, then incubated with the secondary polymer HRP for 1 hour at room temperature. After additional washes, the first Opal fluorophore was incubated with the slides for 10 minutes at room temperature, followed by further washes in TBS-T. This process was repeated from the microwave treatment step for each additional primary antibody, followed by one final repetition of the microwave treatment to strip the primary-secondary antibody complex from the tissue. Antibodies, concentrations and relevant Opal fluorophores can be found in Table S4. Once all primary antibodies had been introduced, slides were counterstained with DAPI for 5 minutes at room temperature, washed with TBS-T and coverslips were mounted using ProLong Diamond Antifade mounting reagent (ThermoFisher Scientific). Multispectral imaging was carried out using the Vectra Quantitative Pathology Imaging system, applying quantitative unmixing of fluorophores and removal of tissue autofluorescence. Images were visualized using the HALO image analysis platform (Indica Labs).

## Statistical analysis

RNA-seq count and PSI data were analyzed as described above. Enrichment of SFs/RBPs in specific clusters was determined by Fisher’s exact test in R. Western blot protein bands were quantified by densitometry analysis in ImageJ and normalized to GAPDH for each sample, and the resulting values were subjected to unpaired student’s t-test. For tau isoform analysis, ratios between each isoform were calculated per sample prior to statistical analysis. For assessment of RNA pull-downs, the amount of eluted SF/RBP protein was normalized to the total amount of SF/RBP protein (Flow-through + Eluate) to determine the percentage of SF/RBP protein bound to the labelled RNA, and this value was used for statistical analysis by student’s t-test. For ISO-seq data, the expression of each isoform was calculated as a proportion of all detected isoforms, and average expression between control, AD and PSP cases was calculated. Statistical difference in isoform expression and expression ratios were calculated by one-way ANOVA with Tukey post-hoc testing. qRTPCR gene expression was analyzed using the ΔΔCt method, and expression was normalized to β-actin as endogenous controls. Statistical significance was determined by the appropriate one-way ANOVA and Bonferroni post-hoc testing. For cell culture experiments, all tests were conducted in triplicate in three independent experiments (total replicates = 9). For human brain analyses, tissue was acquired for 4-6 PSP cases, 4-6 AD cases and 4-6 healthy aged controls. Significant comparisons are labelled in figures as \**p* < 0.05, \*\**p* < 0.01 and \*\*\**p* < 0.001.

## ACKNOWLEDGEMENTS

This work was supported by funding from the BrightFocus Foundation (KRB), Association for Frontotemporal Degeneration (KRB), CurePSP (KRB), the Rainwater Charitable Foundation (AMG, KRB), and Alzheimer’s Association AARF-17-529888 (JDC). We thank the NIH Neurobiobanks at the University of Maryland, Harvard and Mount Sinai for supplying tissues for analyses. We also thank the Neuropathology brain bank and research CoRE and the Genetics CoRE at the Icahn School of Medicine at Mt. Sinai for supplying tissues and conducting immunohistochemical staining, and for carrying out targeted iso-seq and data analysis. We are grateful to the study participants and their families for their contributions to research.

## AUTHOR CONTRIBUTIONS

Conceptualization: KRB, AMG

Methodology: KRB, DAP, LMO, BMJ, KF, KW, AS, JDC

Validation: KRB, DAP, LMO

Formal analysis: KRB, KF

Investigation: KRB, DAP, LMO

Resources: JDC, TR, KF, KW, JFC, AP, AMG

Data curation: KRB

Writing – Original draft: KRB

Writing – Review and editing: KRB, DAP, LMO, KF, KW, JDC, JFC, AP, AMG

Visualization: KRB

Supervision: AMG

Funding Acquisition: KRB, JFC, AMG

## DECLARATION OF INTERESTS

AMG: Scientific advisory board (SAB) for Denali Therapeutics (2015-2018), SAB for Pfizer (2019), SAB for Genentech, consultant for GSK, AbbVie, Biogen and Eisai. All other authors declare no competing interests.

## SUPPLEMENTARY INFORMATION

### Supplementary Figure Legends

**Figure S1.**
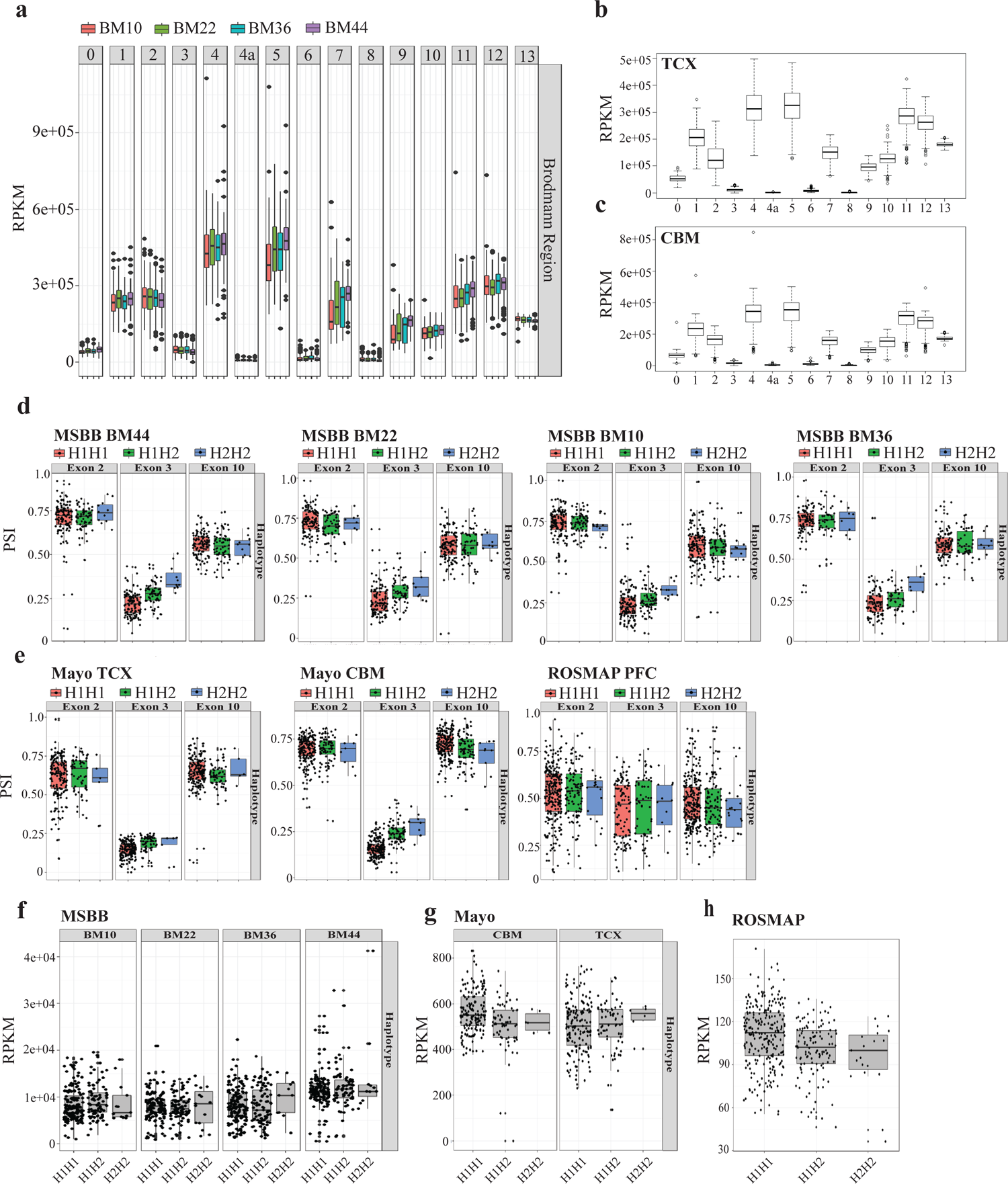
*MAPT* exon 3 inclusion varies between 17q21.31 haplotypes in multiple brain regions. Related to Figure 1. **A)** *MAPT* exon expression (RPKM) for each AMP-AD MSBB brain region. **B-C)** *MAPT* exon expression (RPKM) in temporal cortex (TCX, ***B.***) and cerebellum (CBM, ***C***.) in the AMP-AD MAYO cohort. **D)** PSI values for *MAPT* exons 2, 3 and 10 for each MSBB brain region, split by 17q21.31 haplotype. **E)** PSI values for *MAPT* exons 2, 3 and 10 for both MAYO brain regions and ROSMAP PFC, split by 17q21.31 haplotype. **F-H)** Total *MAPT* expression (RPKM) for each 17q21.31 haplotype in ***F***. MSBB brain regions, ***G***. MAYO brain regions and ***H***. ROSMAP data. All error bars ± SEM

**Figure S2.**
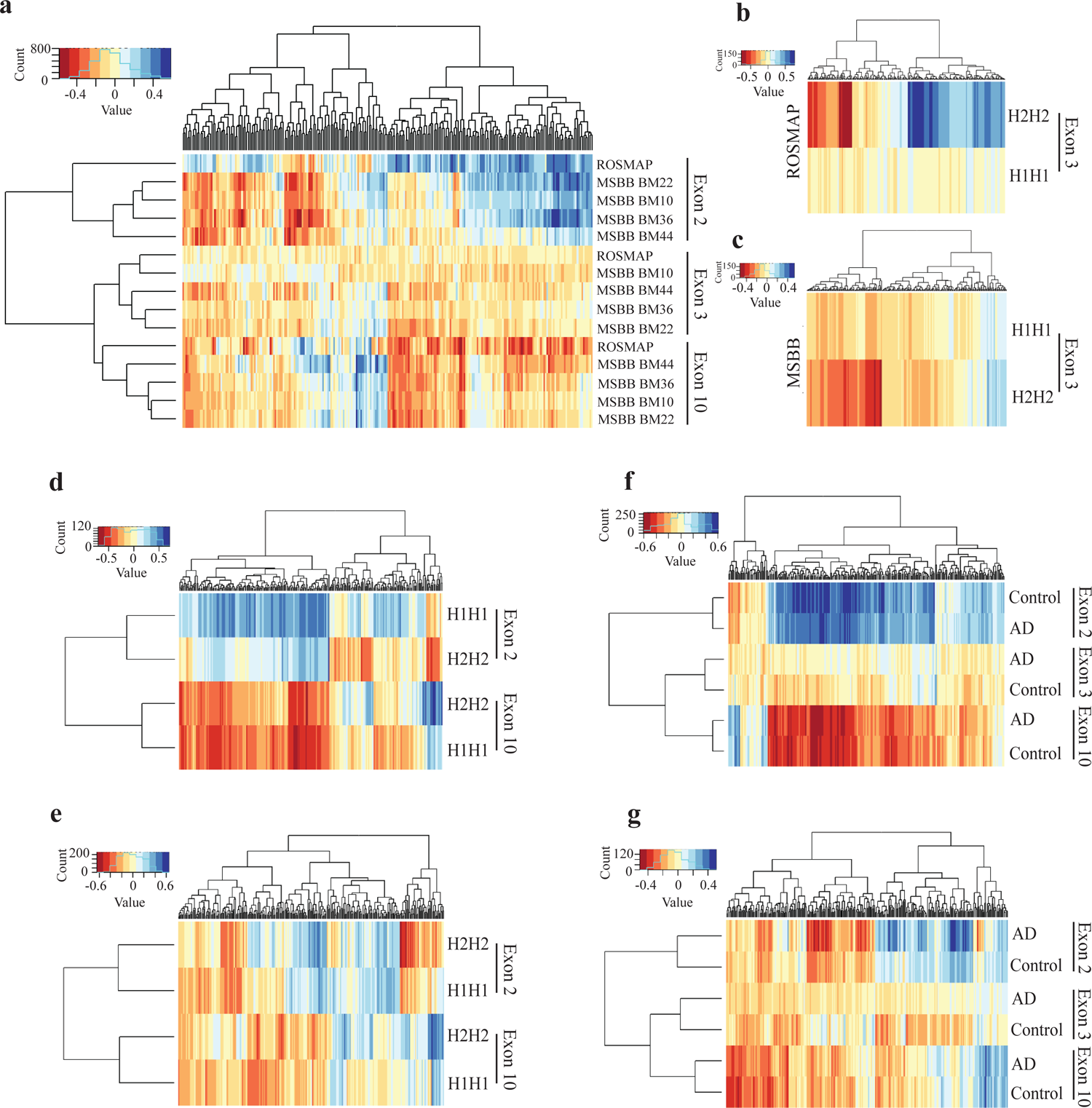
SF/RBP correlations with *MAPT* exons 2 and 10 inclusion are highly correlated between haplotypes and disease status. Related to Figure 1. **A)** Pearson’s correlation coefficients between SF/RBP expression and *MAPT* exon 2, 3 and 10 PSI values following unsupervised hierarchical clustering in ROSMAP and MSBB data. **B-C)** Pearson’s correlation coefficients between SF/RBP expression and *MAPT* exon 3 PSI split between *MAPT* 17q21.31 H1H1 an H2H2 haplotypes in ***B***. ROSMAP and ***C***. MSBB data. **D-E)** Pearson’s correlation coefficients between SF/RBP expression and *MAPT* exon 2 and exon 10 PSI values split between *MAPT* 17q21.31 H1H1 an H2H2 haplotypes in ***D***. ROSMAP and ***E.*** MSBB data. **F-G)** Pearson’s correlation coefficients between SF/RBP expression and *MAPT* exons 2, 3 and 10 PSI values, split between AD and control diagnosis in ***F***. ROSMAP and ***G.*** MSBB data.

**Figure S3.**
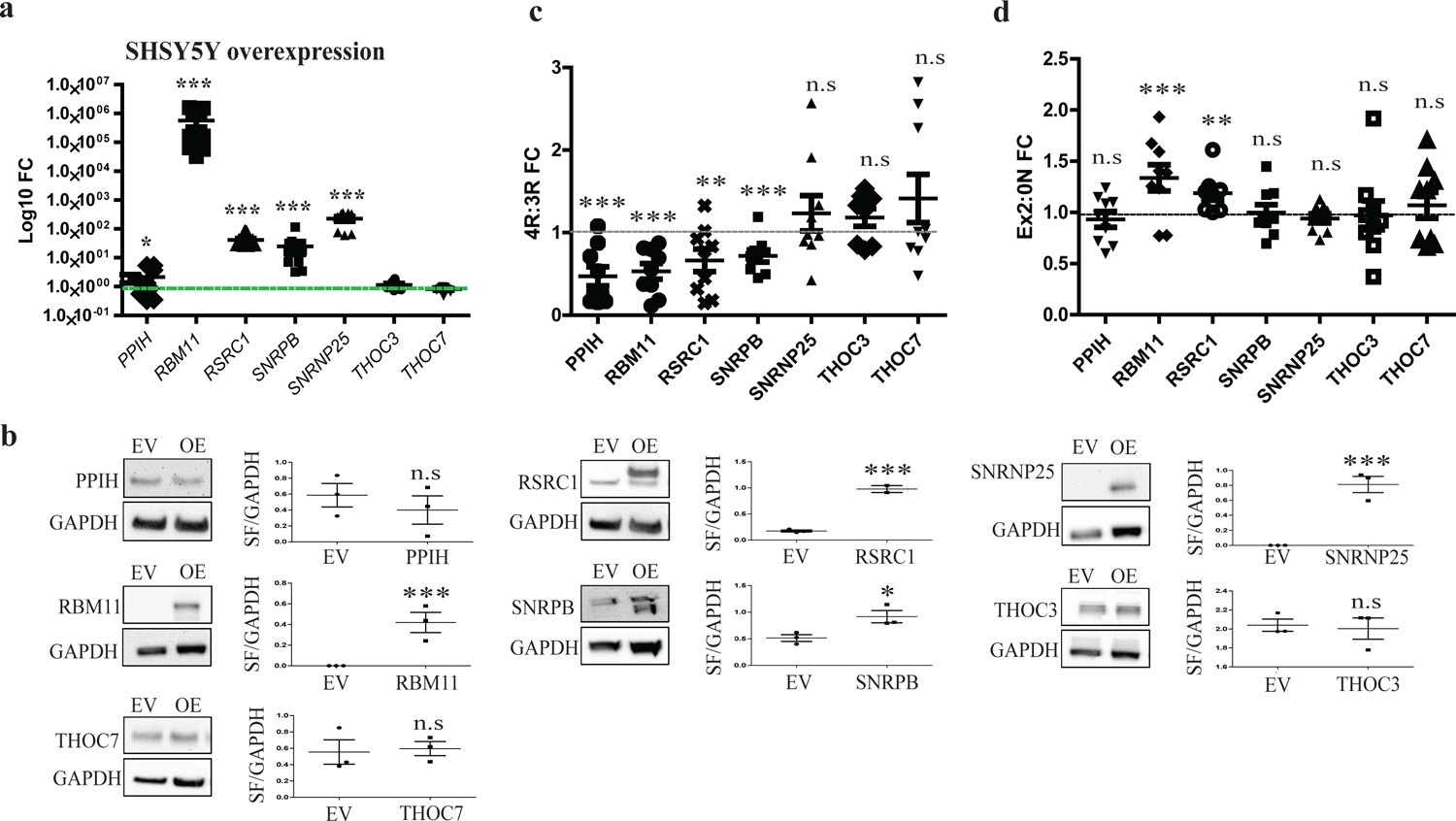
SF/RBP overexpression influences *MAPT* splicing. Related to Figure 2. **A)** Log10 fold change (FC) of each SF/RBP following overexpression in SH-SY5Y cells, normalized to *ACTB* endogenous control. Green line indicates average expression of empty vector control. N = 3 from 3 independent experiments. Student’s t-test \**p* < 0.05, \*\*\**p* < 0.001. **B)** Representative images of western blot validation of SF/RBP overexpression in SH-SY5Y cells with densitometry quantification, normalized to GAPDH (SF/GAPDH). EV = Empty vector control, OE = SF/RBP overexpression. N = 3, Student’s t-test \**p* < 0.05, \*\*\**p* < 0.001, n.s = not significant **C-D)** Expression fold change (FC) of the ***C***. 4R:3R ratio and ***D***. Exon2:0N ratio in SH-SY5Y cells following SF/RBP overexpression. Grey lines indicate average expression of empty vector control. N = 3 from 3 independent experiments. Student’s t-test \**p* < 0.05, \*\**p* < 0.01, \*\*\**p* < 0.001, n.s = not significant.

**Figure S4.**
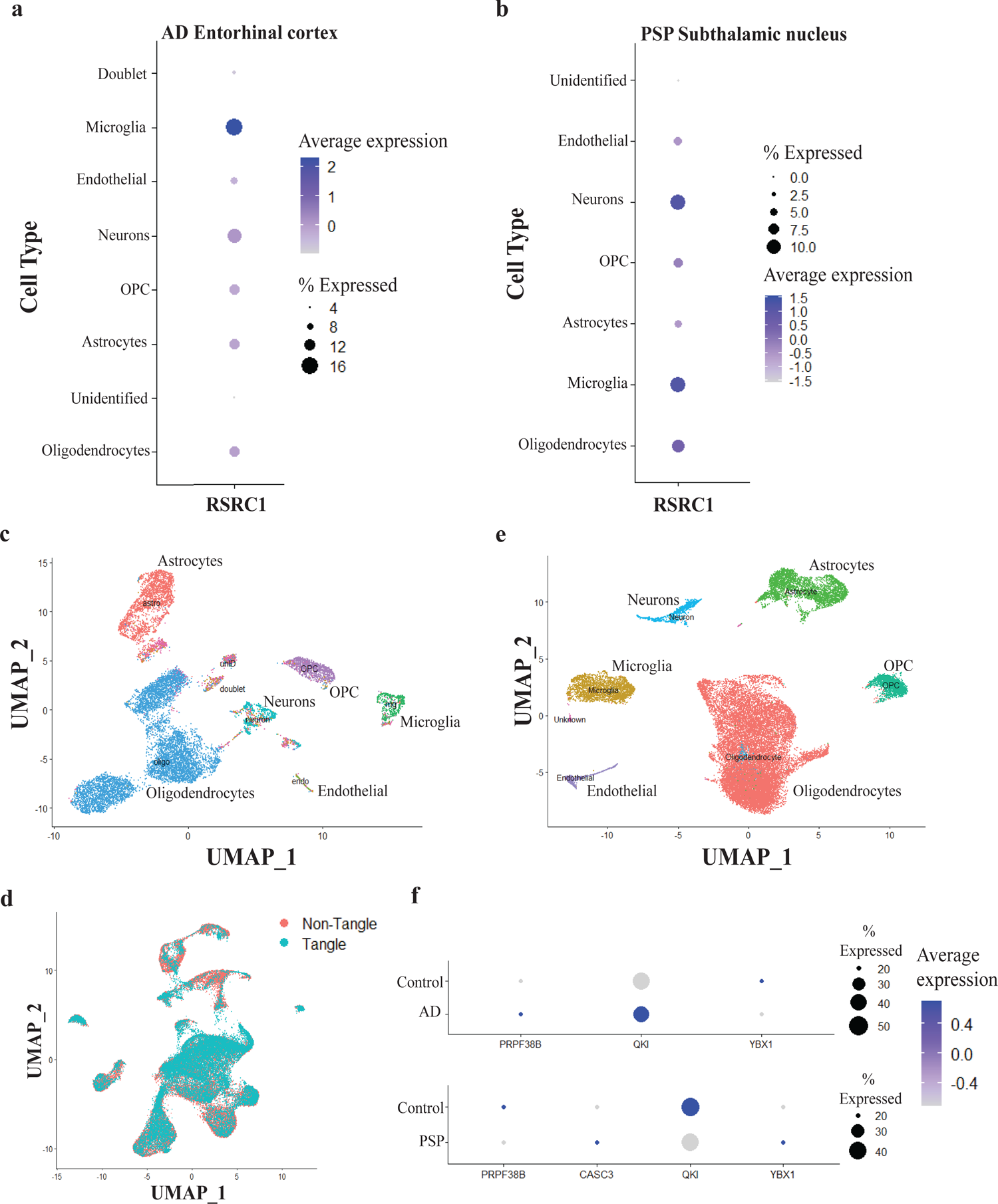
*RSRC1* is expressed in multiple neural cell types, including neurons. Related to Figure 3. **A-B)** Expression of *RSRC1* in neural cell types detected by snuc-seq in ***A.*** entorhinal cortex and ***B.*** subthalamic nucleus. Average expression is scaled across cell types, and deeper colors represent higher gene expression. Dot size indicates proportion of *RSRC1* expressing cells. **C)** UMAP reduction of snuc-seq data from AD and control entorhinal cortex, with clusters colored by cell type, as defined in Grubman *et al*. 2019. **D)** UMAP reduction of single-soma data from AD prefrontal cortex, colored by AT8 positive (“Tangle”) or AT8 negative (“Non-Tangle”) neurons. **E)** UMAP reduction of snuc-seq data from PSP and control subthalamic nucleus, with clusters colored by cell type, as defined by positivity for markers described in Grubman *et al.* 2019. **F)** Expression of *MAPT* exon 2 includer genes in AD (top) and PSP (bottom) neurons from snuc-seq data. Dot size represents the proportion of neurons expressing the gene, depth of color indicates normalized average gene expression.

**Figure S5.**
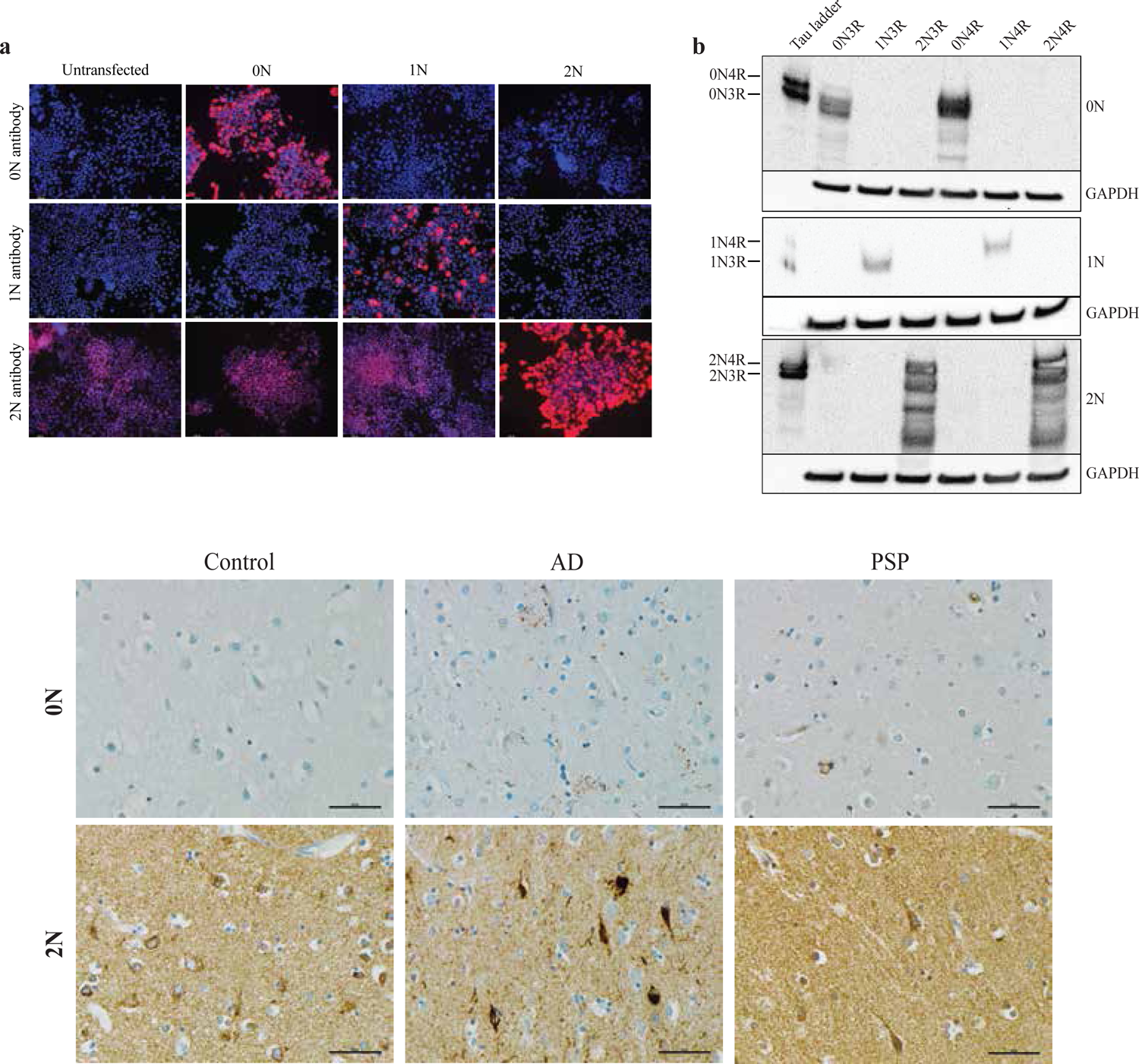
N-terminal tau antibodies are specific for each tau isoform. Related to Figure 4. **A)** N2a cells overexpressing either 0N3R (0N), 1N3R (1N), 2N3R (2N) or untransfected controls, labelled with Abcam 0N and 2N tau antibodies, and BioLegend 1N tau antibody. **B)** Western blot of N2a cells overexpressing each tau isoform, detected by 0N, 1N and 2N tau N-terminal antibodies. Band size compared to tau ladder. GAPDH used as a loading control. **C)** IHC detection of N-terminal tau in control, AD and PSP temporal cortex using alternative antibodies (Abcam 0N, 2N) to those in Figure 4 (BioLegend 0N, 2N).

**Figure S6.**
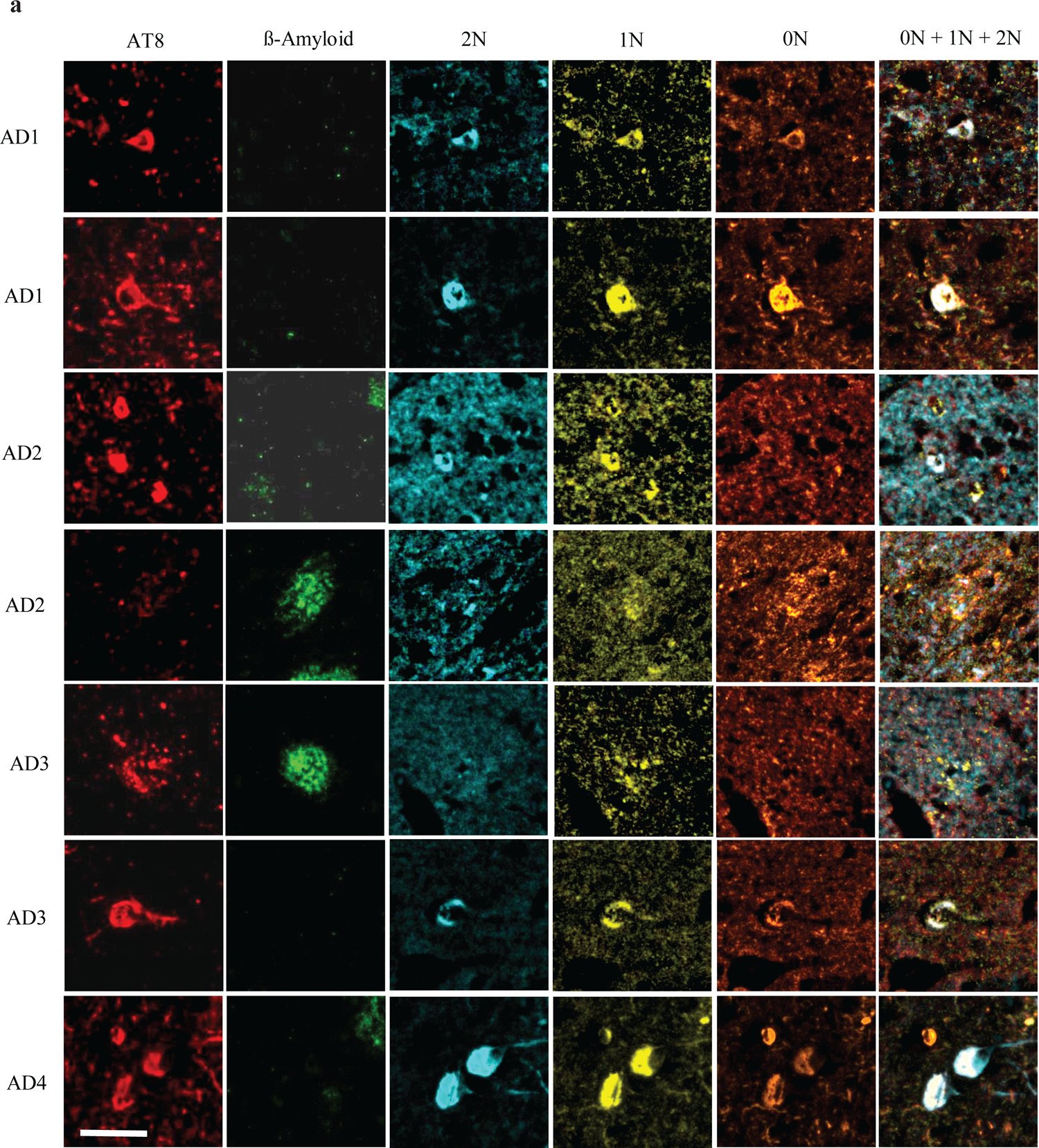
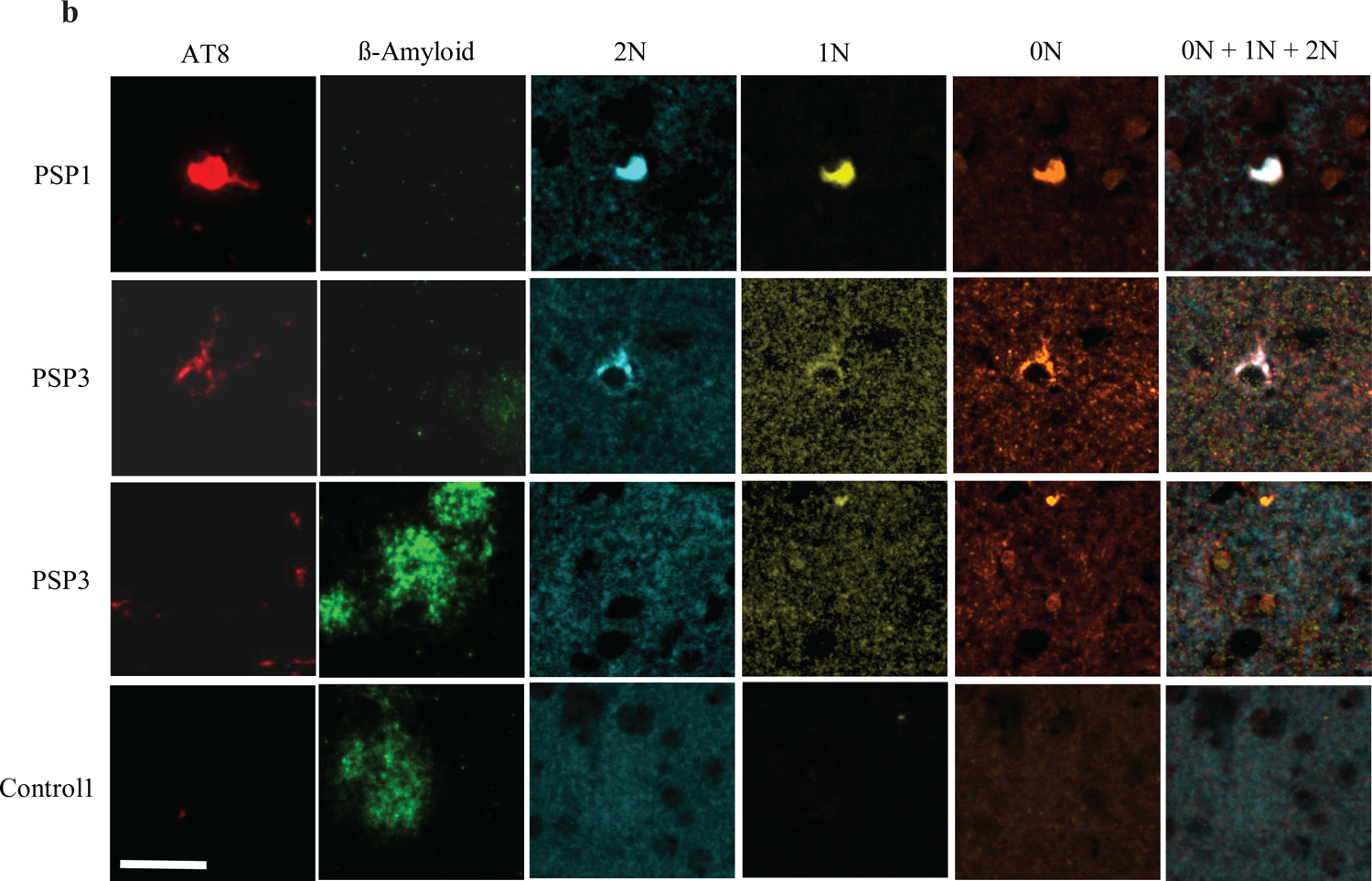
N-terminal tau accumulates in AD and PSP brain. Related to Figure 5. **A)** Representative images of multiplex immunofluorescent labeling of AD temporal cortex with AT8 (red), β-amyloid (green), 2N tau (blue), 1N tau (yellow) and 0N tau (orange), and overlay of all three N-terminal tau antibodies in 4 different individuals. Examples of tau accumulation in neurofibrillary tangles (AD1-4) and dystrophic neurites surrounding amyloid plaques (AD2-3). N=4, scale bar = 50µm. **B)** Representative images of immunofluorescent labeling of PSP and control temporal cortex as in ***A.*** Examples of neurofibrillary tangles (PSP1, 3) and absence of tau-positive dystrophic neurites surrounding amyloid plaques in PSP (PSP3) and control brain (Control1). N=4, scale bar = 50µm

## Supplementary tables

**Table S1.**
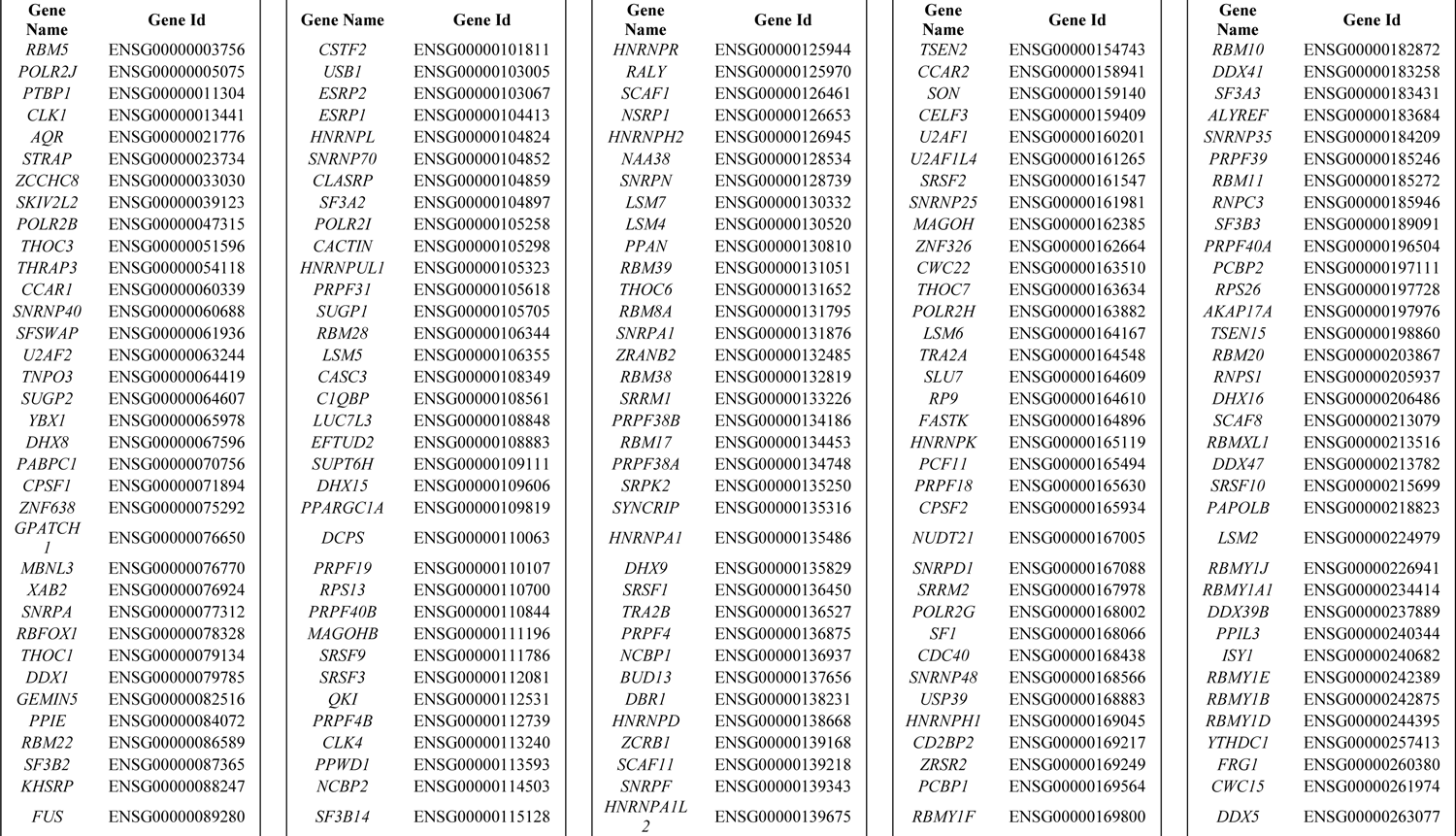
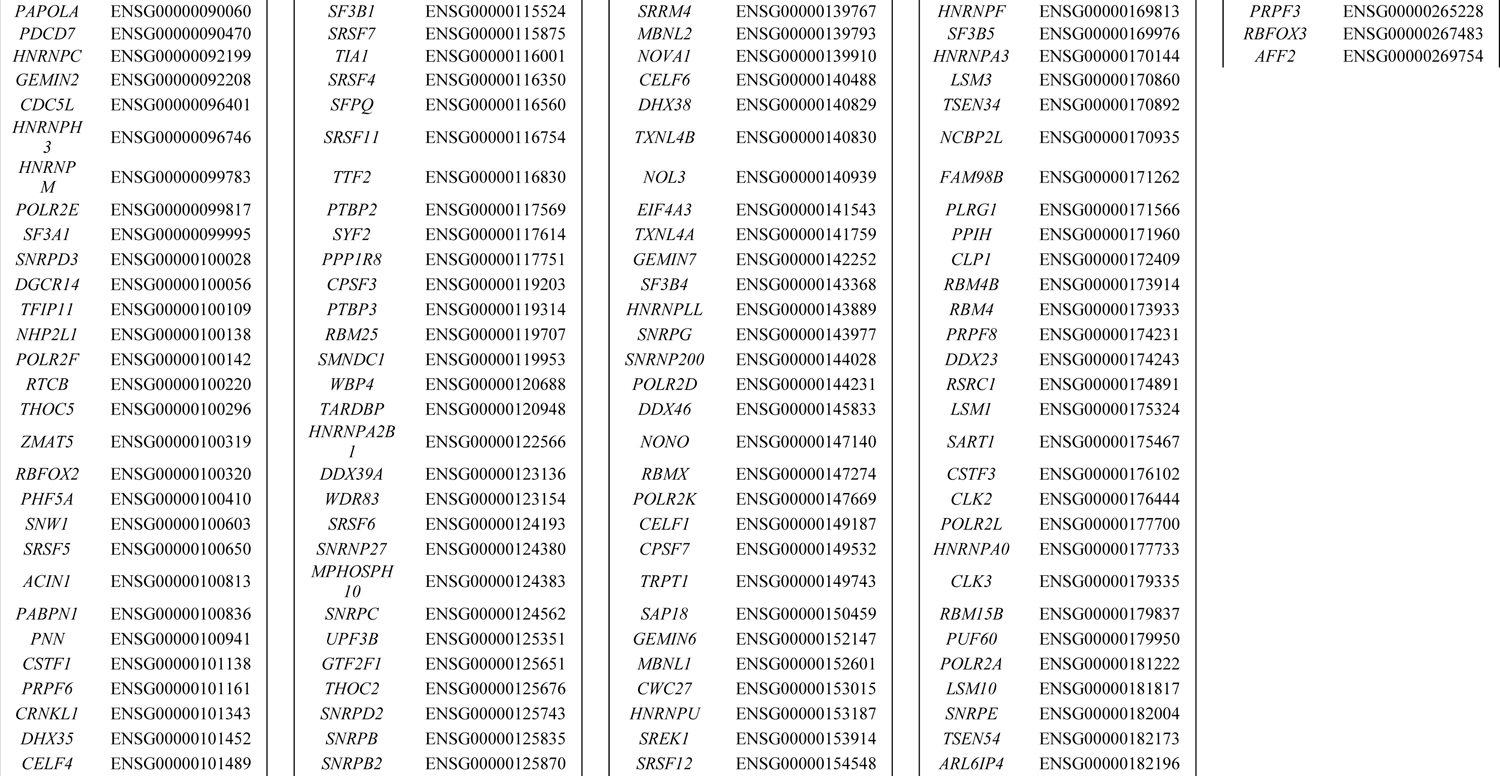
SF/RBP genes included in MAPT exon PSI analyses

**Table S2.**
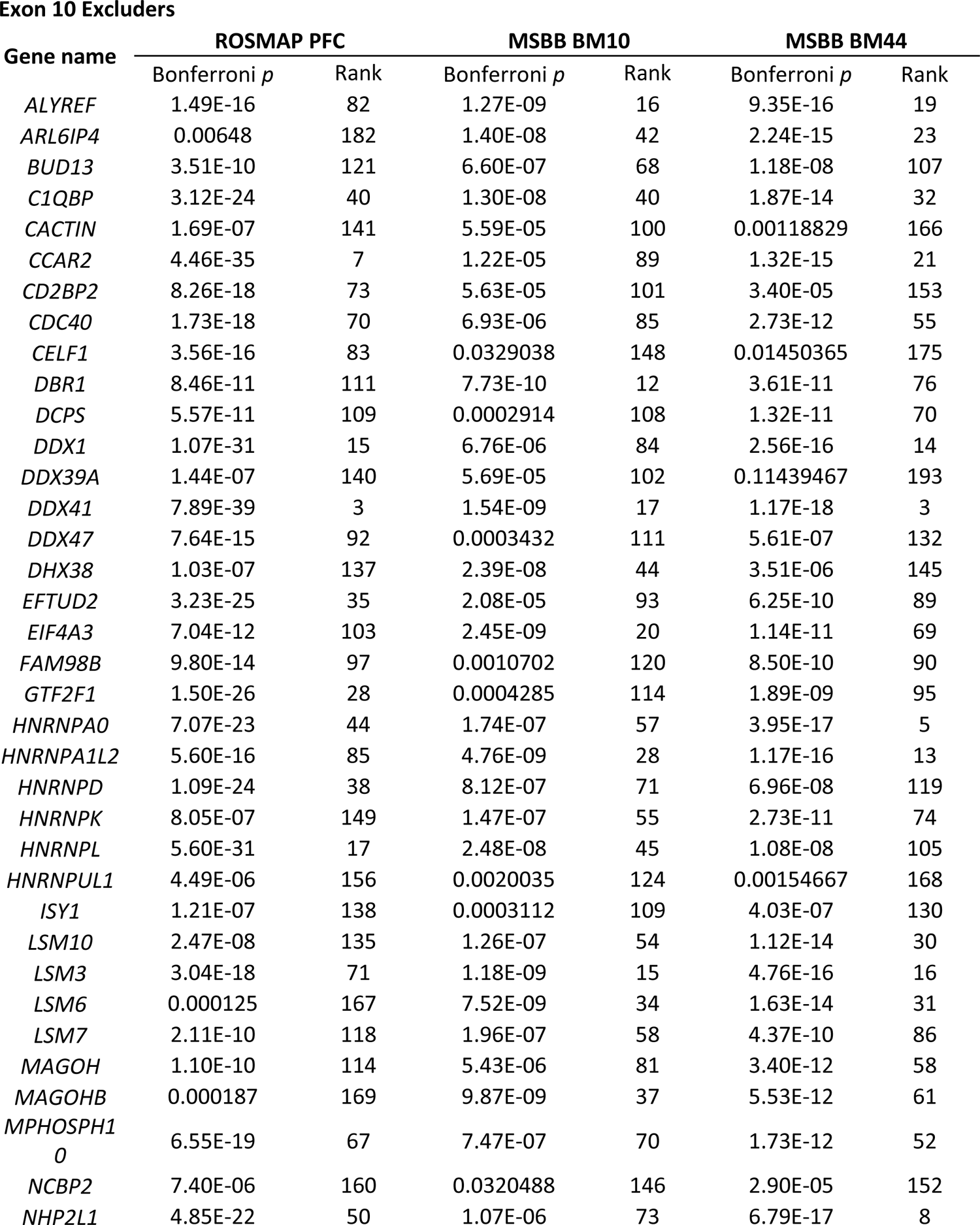
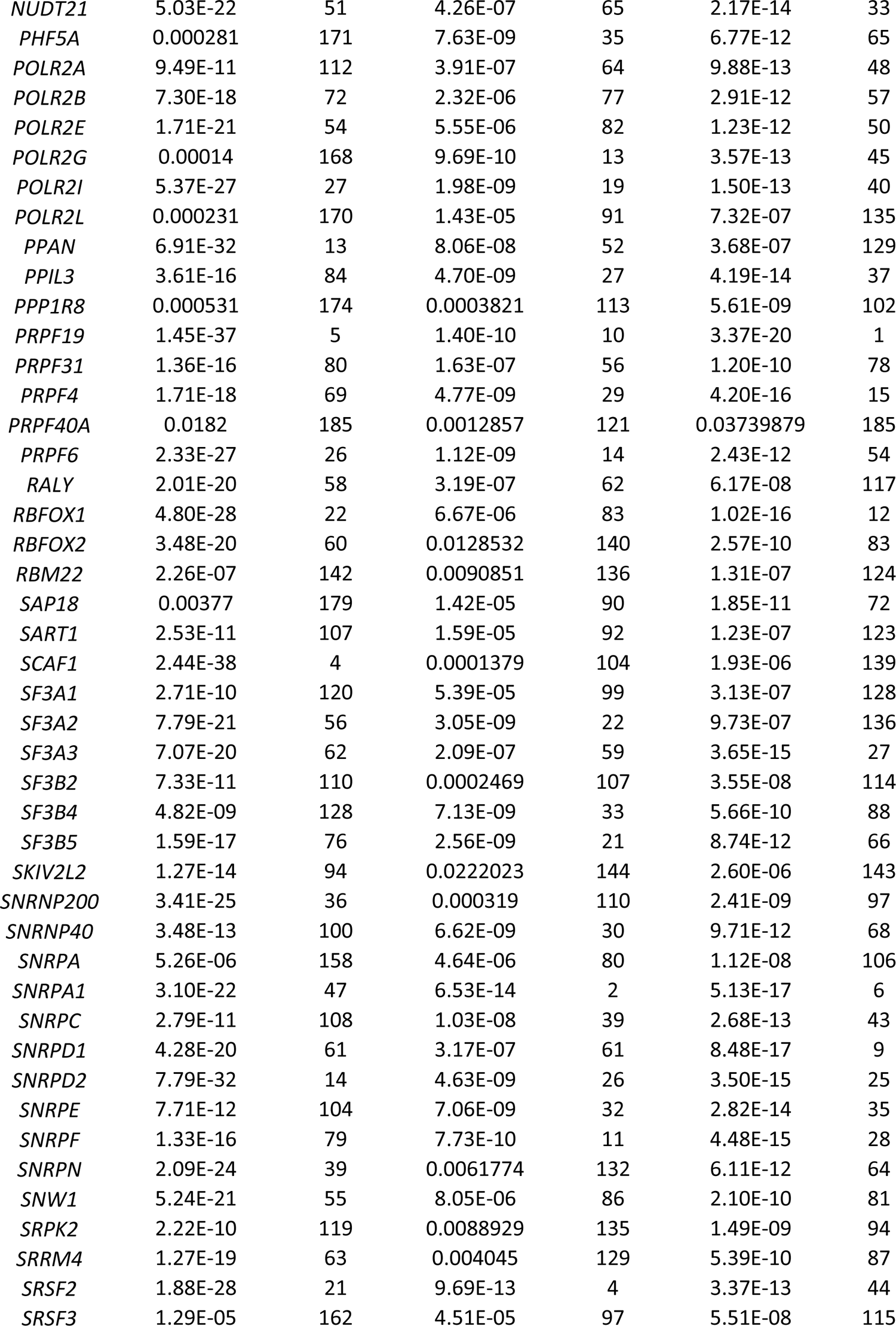
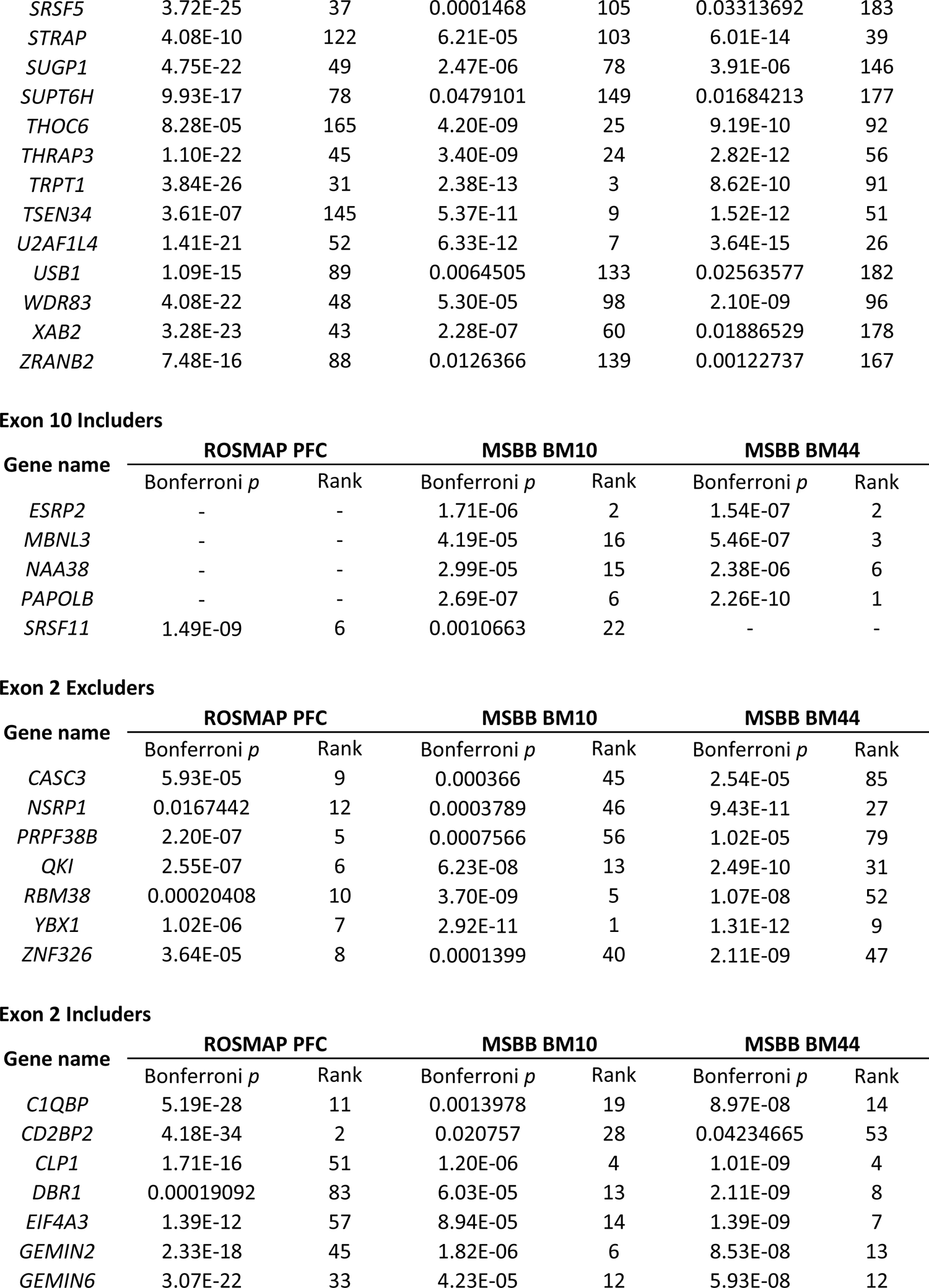
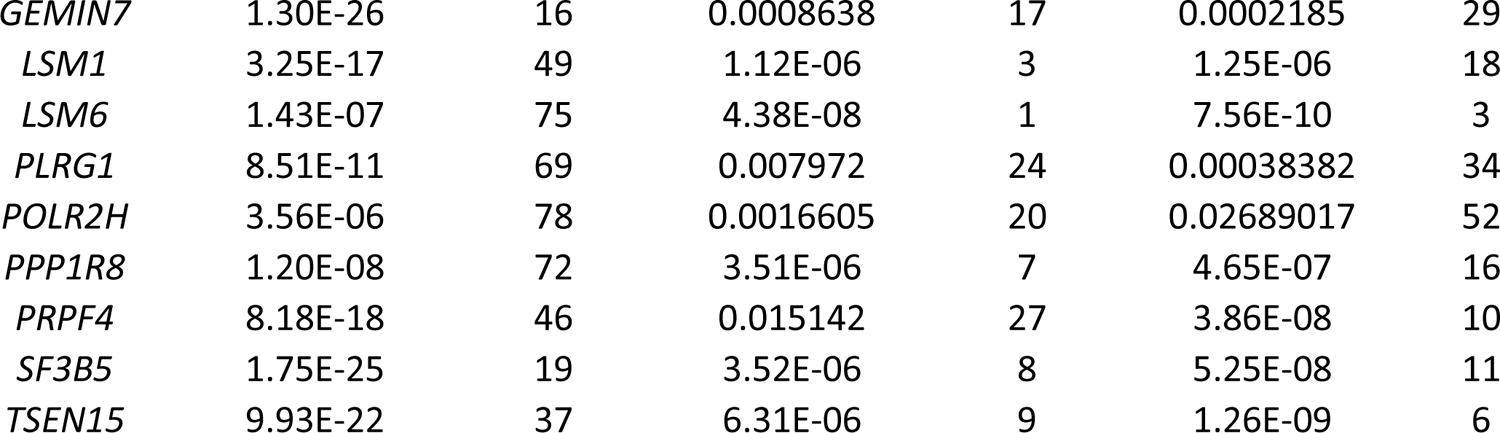
SF/RBPs significantly associated with *MAPT* exon 2 or 10 splicing in ROSMAP, MSBB BM10 and MSBB BM44 RNA-seq data

**Table S3.** Summary of human brain tissues and sources used in this study

| ID | Dx | AoD | Sex | Source |
| --- | --- | --- | --- | --- |
| **Western blot & ISO-seq** |  |  |  |  |
| 9 | Contro<br>l | 55 | Male | ISMMS Neuropathology Research Core and Brain Bank |
| 10 | Contro<br>l | 63 | Male | ISMMS Neuropathology Research Core and Brain Bank |
| 11 | Contro<br>l | 73 | Male | ISMMS Neuropathology Research Core and Brain Bank |
| 13 | Contro<br>l | 74 | Female | ISMMS Neuropathology Research Core and Brain Bank |
| 19 | Contro<br>l | 18 | Male | ISMMS Neuropathology Research Core and Brain Bank |
| 22 | Contro<br>l | 72 | Female | ISMMS Neuropathology Research Core and Brain Bank |
| S08611 | AD | 81 | Male | NIH Neurobiobank - Harvard |
| 38917 | AD | 90 | Female | NIH Neurobiobank - Mt.Sinai |
| S13471 | AD | 86 | Female | NIH Neurobiobank - Harvard |
| S15985 | AD | 85 | Female | NIH Neurobiobank - Harvard |
| S05093 | AD | 85 | Male | NIH Neurobiobank - Harvard |
| 3853 | AD | 85 | Male | NIH Neurobiobank - Harvard |
| 5737 | PSP | 73 | Male | NIH Neurobiobank - Maryland |
| 6191 | PSP | 62 | Female | NIH Neurobiobank - Maryland |
| 5126 | PSP | 72 | Male | NIH Neurobiobank - Maryland |
| 6043 | PSP | 67 | Female | NIH Neurobiobank - Maryland |
| 6225 | PSP | 87 | Male | NIH Neurobiobank - Maryland |
| 6085 | PSP | 88 | Female | NIH Neurobiobank - Maryland |
| **IHC & IF** |  |  |  |  |
| Control1 | Contro<br>l | 73 | Male | ISMMS Neuropathology Research Core and Brain Bank |
| Control2 | Contro<br>l | 68 | Female | ISMMS Neuropathology Research Core and Brain Bank |
| Control3 | Contro<br>l | 66 | Female | ISMMS Neuropathology Research Core and Brain Bank |
| AD1 | AD | 85 | Female | ISMMS Neuropathology Research Core and Brain Bank |
| AD2 | AD | 95 | Male | ISMMS Neuropathology Research Core and Brain Bank |
| AD3 | AD | 78 | Female | ISMMS Neuropathology Research Core and Brain Bank |
| AD4 | AD | 66 | Female | ISMMS Neuropathology Research Core and Brain Bank |
| PSP1 | PSP | 64 | Male | ISMMS Neuropathology Research Core and Brain Bank |
| PSP2 | PSP | 71 | Male | ISMMS Neuropathology Research Core and Brain Bank |
| PSP3 | PSP | 69 | Male | ISMMS Neuropathology Research Core and Brain Bank |

***Dx = Diagnosis. AoD = Age of death.***

**Table S4.**
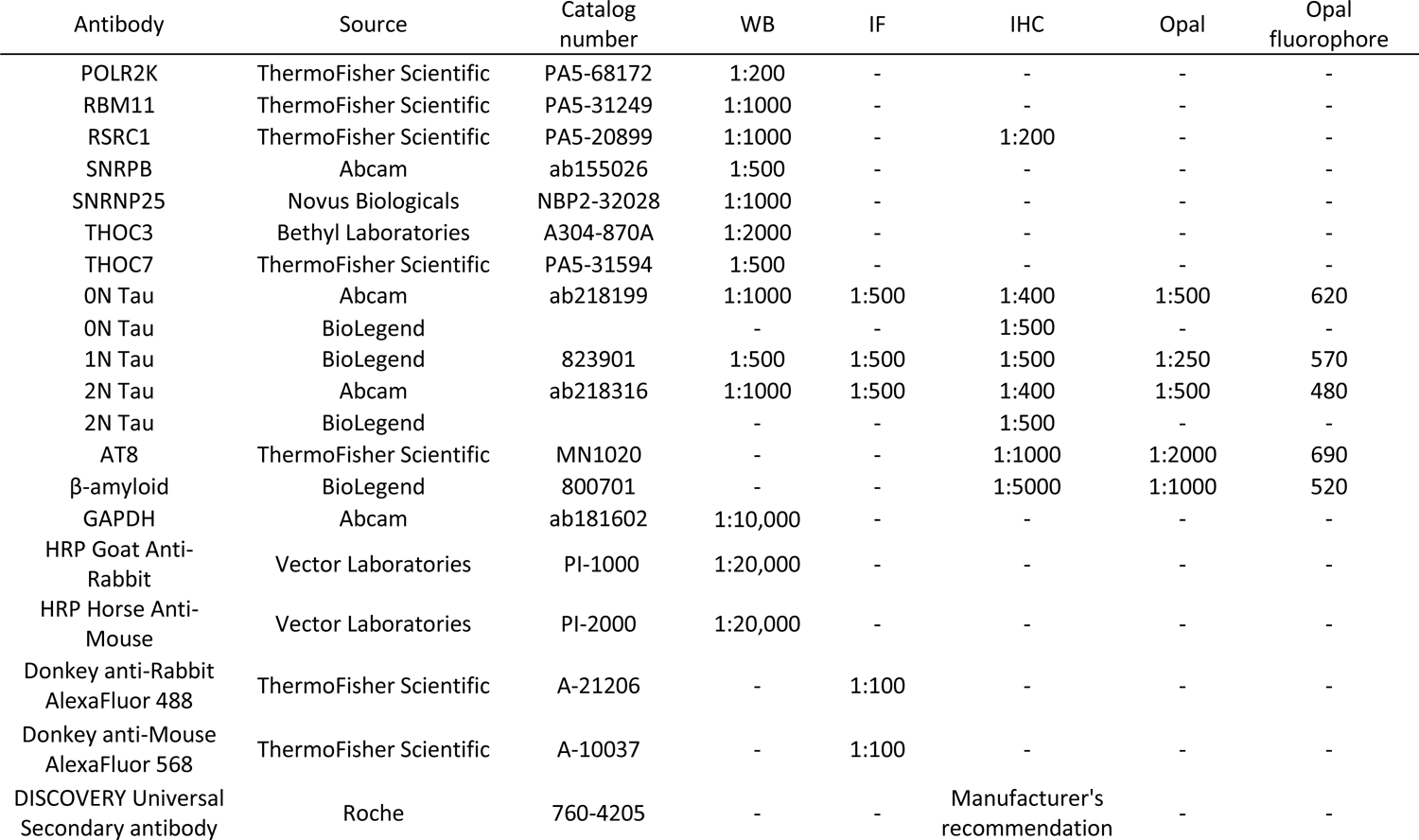
Summary of all antibodies and dilutions used in this study

| Antibody | Source | Catalog number | WB | IF | IHC | Opal | Opal fluorophore |
| --- | --- | --- | --- | --- | --- | --- | --- |
| POLR2K | ThermoFisher Scientific | PA5-68172 | 1:200 | - | - | - | - |
| RBM11 | ThermoFisher Scientific | PA5-31249 | 1:1000 | - | - | - | - |
| RSRC1 | ThermoFisher Scientific | PA5-20899 | 1:1000 | - | 1:200 | - | - |
| SNRPB | Abcam | ab155026 | 1:500 | - | - | - | - |
| SNRNP25 | Novus Biologicals | NBP2-32028 | 1:1000 | - | - | - | - |
| THOC3 | Bethyl Laboratories | A304-870A | 1:2000 | - | - | - | - |
| THOC7 | ThermoFisher Scientific | PA5-31594 | 1:500 | - | - | - | - |
| 0N Tau | Abcam | ab218199 | 1:1000 | 1:500 | 1:400 | 1:500 | 620 |
| 0N Tau | BioLegend |  | - | - | 1:500 | - | - |
| 1N Tau | BioLegend | 823901 | 1:500 | 1:500 | 1:500 | 1:250 | 570 |
| 2N Tau | Abcam | ab218316 | 1:1000 | 1:500 | 1:400 | 1:500 | 480 |
| 2N Tau | BioLegend |  | - | - | 1:500 | - | - |
| AT8 | ThermoFisher Scientific | MN1020 | - | - | 1:1000 | 1:2000 | 690 |
| β-amyloid | BioLegend | 800701 | - | - | 1:5000 | 1:1000 | 520 |
| GAPDH | Abcam | ab181602 | 1:10,000 | - | - | - | - |
| HRP Goat Anti-Rabbit | Vector Laboratories | PI-1000 | 1:20,000 | - | - | - | - |
| HRP Horse Anti-Mouse | Vector Laboratories | PI-2000 | 1:20,000 | - | - | - | - |
| Donkey anti-Rabbit AlexaFluor 488 | ThermoFisher Scientific | A-21206 | - | 1:100 | - | - | - |
| Donkey anti-Mouse AlexaFluor 568 | ThermoFisher Scientific | A-10037 | - | 1:100 | - | - | - |
| DISCOVERY Universal Secondary antibody | Roche | 760-4205 | - | - | Manufacturer's recommendation | - | - |

## Notes

### Competing Interest Statement

The authors have declared no competing interest.

